# Natural variation in the contribution of microbial density to inducible immune dynamics

**DOI:** 10.1101/788281

**Authors:** Derrick Jent, Abby Perry, Justin Critchlow, Ann T. Tate

**Affiliations:** Department of Biological Sciences, Vanderbilt University. Nashville, TN 37232

**Keywords:** ecological immunology, infection tolerance, life history trade-offs, host-parasite interactions, innate immunity, *Tribolium castaneum*, *Bacillus thuringiensis*

## Abstract

Immune responses evolve to balance the benefits of microbial killing against the costs of autoimmunity and energetic resource use. Models that explore the evolution of optimal immune responses generally include a term for constitutive immunity, or the level of immunological investment prior to microbial exposure, and for inducible immunity, or investment in immune function after microbial challenge. However, studies rarely consider the functional form of inducible immune responses with respect to microbial density, despite the theoretical dependence of immune system evolution on microbe-versus immune-mediated damage to the host. In this study, we analyze antimicrobial peptide (AMP) gene expression from seven wild-caught flour beetle populations (*Tribolium* spp.) during acute infection with the virulent bacteria *Bacillus thuringiensis* (Bt) and *Photorhabdus luminescens* (P.lum) to demonstrate that inducible immune responses mediated by the humoral IMD pathway exhibit natural variation in both microbe density-dependent and independent temporal dynamics. Beetle populations that exhibited greater AMP expression sensitivity to Bt density were also more likely to die from infection, while populations that exhibited higher microbe density-independent AMP expression were more likely to survive *P. luminescens* infection. Reduction in pathway signaling efficiency through RNAi-mediated knockdown of the *imd* gene reduced the magnitude of both microbe-independent and dependent responses and reduced host resistance to Bt growth, but had no net effect on host survival. This study provides a framework for understanding natural variation in the flexibility of investment in inducible immune responses and should inform theory on the contribution of non-equilibrium host-microbe dynamics to immune system evolution.

## Introduction

Mounting and maintaining an immune response is costly to organismal fitness. Investment in immunity requires the diversion of energetic resources away from processes like development and reproduction (Bajgar *et al*. 2015), and immune responses can impose pathological collateral damage upon host tissue (Sadd & Siva-Jothy 2006). In addition, immune defenses that are effective against one parasite might trade off with the production of immune responses effective against other parasites (Murphy *et al*. 2013; Ezenwa *et al*. 2010), or even directly facilitate their colonization (Dejnirattisai *et al*. 2010). These constraints contribute to the conceptual division between constitutive immunity, where hosts invest in immunity prior to exposure to a focal microbe, and inducible immunity, where hosts invest in an immune response, and pay the associated consequences (Ardia *et al*. 2012), only after microbial exposure.

The relative costs and benefits of constitutive and inducible immunity, and the associated implications for immune system evolution, have received appreciable theoretical and experimental attention. For example, an experiment that evolved *Psuedomonas aeruginosa* resistance to a mu-like phage under high and low resource environments found that in high resource environments, the bacteria evolved constitutive resistance to phage through the loss of a surface receptor, while low resource environments favored the inducible CRISPR-Cas system (Westra *et al*. 2015). A mathematical model created by Hamilton and colleagues (Hamilton *et al*. 2008) suggests that highly predictable parasite growth rates should always favor modulation of constitutive investment, whereas variation or unpredictability in parasite growth rates should favor a combination of constitutive and inducible investment. Other models add to these predictions by suggesting that longer inducible time delays (Shudo & Iwasa 2001) and developmental constraints on inducible immune investment (Tate & Graham 2015a) should further favor constitutive defense, while variation in the types of costs associated with constitutive and inducible responses could favor differential investment in recovery rates after infection (Cressler *et al*. 2015).

Inducible immunity is easy to define in broad strokes as the change in immune defense after microbial exposure. However, it is less clear to what extent time and microbe density feed back on its continued production, or how these feedbacks might ultimately influence the costs that drive immune system evolution. The Hamilton *et al*. model (Hamilton *et al*. 2008), for example, assumes that inducible immunity increases exponentially with parasite density, whereas the Shudo and Iwasa model (Shudo & Iwasa 2001) assumes a step-wise rate of inducible response generation that is not sensitive to microbial dynamics. Another model of inducible immune response evolution proposes both microbe density-independent and dependent terms for the rate of immune induction (Frank 2002), but keeps the latter constant when analyzing the sensitivity of host fitness to parameter variation. As with the contrasting cost structure of constitutive and inducible immunity, microbe independent and dependent inducible dynamics should experience trade-offs associated with paying a sunk cost while getting proactive benefits versus investing only when needed but ceding the temporal advantage to the microbes. Intuitively, then, we might predict that microbe-independent inducible responses would be favored against fast-growing, manipulative, or virulent parasites, whereas microbe-dependent terms might be favored by variable parasite exposure rates or virulence characteristics that depend on microbe density. We might additionally expect that microbe density-dependent inducible immunity would be useful in controlling endosymbiont or gut microbe populations (Login *et al*. 2011), where host fitness and the benefits of tolerance may be optimized at intermediate microbe density, disfavoring systems that are switched on by the mere presence or absence of microbes.

A better understanding of natural variation in the sensitivity of inducible immune dynamics to time and changes in the microbial population, therefore, would improve our conceptual and theoretical frameworks for immune system evolution. Unfortunately, data on the relative rates of microbe dependent and independent inducible immune dynamics are sparser than the mathematical models that employ them. Nevertheless, recent data from fruit flies infected with bacteria (Louie *et al*. 2016) and viruses (Gupta & Vale 2017), as well as flour beetles infected with bacteria (Tate & Graham 2017), suggest that the inducible expression of antimicrobial peptides (AMPs) controlled by humoral signaling pathways (particularly IMD, but also Toll and Jak-STAT) are strongly correlated to microbe density during both acute (Tate & Graham 2017) and recovery (Louie *et al*. 2016) phases of infection. These studies suggest that microbe density-dependent feedbacks might indeed be an important component of inducible dynamics. However, all these studies were performed using a few laboratory host strains. A better understanding of natural variation in the functional form of inducible dynamics with respect to microbe density would inform future mathematical models of immune system evolution and promote deeper consideration of a ubiquitous immune system trait.

To test the hypothesis that the evolutionary costs and benefits of microbe-independent and – dependent inducible immune responses depend on the context of infection and are thus likely to vary among populations, we infected seven wild-caught flour beetle (*Tribolium castaneum* and *T. confusum*) populations with one of two entomopathogenic bacteria, the rapidly growing and virulent *Bacillus thuringiensis* (Bt) and the slower-growing but immuno-modulatory *Photorhabdus luminescens* (P.lum). To quantify inducible immune gene expression, we infected individuals from each population with a gradient of initial bacterial doses or a sterile wound and then sacrificed them during a point in the acute infection phase that is late enough to avoid lag in the induction of immunity but prior to the onset of host mortality (Tate *et al*. 2017). Using RT-qPCR, we quantify the impact of natural variation and humoral (IMD) immune pathway signaling efficiency on the intercept (microbe-independent) and slope (microbe-sensitive) of the relationship between microbe density and immune gene transcription. We then identify intriguing correlations between immune parameters and host resistance and survival against bacterial infection. These results give basic insight into the evolution and dynamics of inducible immune responses, and should inspire future experiments that assay microbe density so that we can better understand the costs and benefits of immune sensitivity to microbial burden.

## Materials and Methods

### Beetle sources and rearing

The wild-derived beetles used in these experiments were obtained by setting pheromone-baited dome traps (Trece) at feed stores and grain elevators around Pennsylvania (June 2013; Snavely *T. confusum* and *T. castaneum*), middle Tennessee (June 2017; Marshall *T. confusum*) and southern Kentucky (June 2017; Green River *T. confusum* and *T. castaneum*, Dorris *T. castaneum*, WF Ware *T. castaneum*). Traps were reset and beetles collected weekly for five weeks. We created breeding populations in the lab from at least 20 active adult beetles per population. Most populations were naturally infected with protozoan parasites, so to create uninfected colonies, we bathed eggs derived from each breeding population in 1% Virkon and rinsed them 2 times in saline solution before passaging them to clean flour for hatching. This method reliably produces protozoan-free populations (Tate & Graham 2015b), and lack of parasites was confirmed by dissection of 20 adult and larval beetles in the F1 and F2 generations coupled with inspection of frass via microscopy for presence of gametocysts. All beetle populations were also inspected for infection with other bacteria, protozoa, and microsporidia via dissection and microscropy, and were observed for 2 generations to ensure no unexplained mortality.

### Creation of experimental groups

For all experiments, we derived experimental larvae from breeding groups created by placing 40 adults from a stock colony in a petri dish with flour and allowing them to lay eggs for 48 hours, whereupon the adults were passaged to a new petri dish and larvae were provided with *ad libitum* flour through development. As size can influence mortality rates, 4.5mm larvae were selected from these breeding groups for infection experiments, thus controlling for both size and age.

### Bt Infection Experiments

*Bacillus thuringiensis* is a spore-producing entomopathogen that relies on host killing for transmission (Raymond *et al*. 2010). Whether it naturally enters the host through the gut or is septically injected, it grows rapidly in the hemolymph of infected beetles and kills them within 12 hours (Tate *et al*. 2017). To infect beetles with Bt (ATCC 55177, a berliner strain that contains a cry protein strain active against Coleoptera), we prepared a culture from a glycerol stock kept at −80°C and grown overnight in 10mL of Nutrient Broth #3 (Sigma Aldrich). In the morning, we transferred 200uL of the overnight culture to 3 mL of fresh NB and brought to an OD of 0.5 to produce the log phase culture. We created the initial doses for all experiments by mixing 500uL overnight culture with 500uL log phase culture (approximately 1*10^8^ colony forming units (CFU)/mL), and then aliquoting a volume of this mixture to sterile NB with volumes that produced four two-fold dilutions (dose 1 (∼50 CFU/needle prick), dose 2 (100 CFU, the dose that kills ∼50% of hosts (LD50)), dose 4 (200 CFU), and dose 8 (800 CFU)), as well as a sterile NB control (dose “0”) or a naïve control where insects were handled but not pricked. We infected beetles by dipping an ultrafine insect pin in the bacterial or injection control solutions, and then punctured the beetle under the sixth segment (larvae) or between the head and pronotum (adults). All beetles were kept in individual wells of 96 well plates at 30°C after treatment.

#### Experiment 1: Bt density and gene expression over 4 initial doses

For bacterial and gene expression assays, all beetles from each population (N = 6-10 beetles/dose/population) were sacrificed at 8 hours post infection by flash freezing and stored at −80°C until processing. We stratified beetles into different bacterial dose treatments, as described above, to ensure a continuous gradient of bacterial density spanning several orders of magnitude upon cross-sectional sampling.

#### Experiment 2: Host survival after Bt infection and bacterial load at time of death

For survival assays (N = 50-60/population/experiment), individual mortality after infection with dose 2 (100 CFU/needle prick) or sterile NB (wounding control) was monitored every 30 minutes for 12 hours (peak mortality 8-11 hours) and then again at 24 hours post infection. Survival experiments were conducted in two blocks. The first included all populations except for Marshall, which was inadvertently excluded from breeding group setup, and the second included Marshall alongside Snavely *T. confusum* and *T. castaneum* as comparison groups. The second block received a slightly higher initial dose (190 CFU/needle prick), resulting in a higher average mortality rate (**Fig. 1A, B**). Individuals that died prior to 4 hours post infection (0-5%) were discarded from the analysis, as mortality is a result of trauma rather than bacterial infection. For estimating bacterial load at time of death (BLUD), individuals were collected as soon as they demonstrated moribund behavior (legs in air, feeble response to touching) and immediately flash-frozen for storage at −80°C.

**Figure 1.**
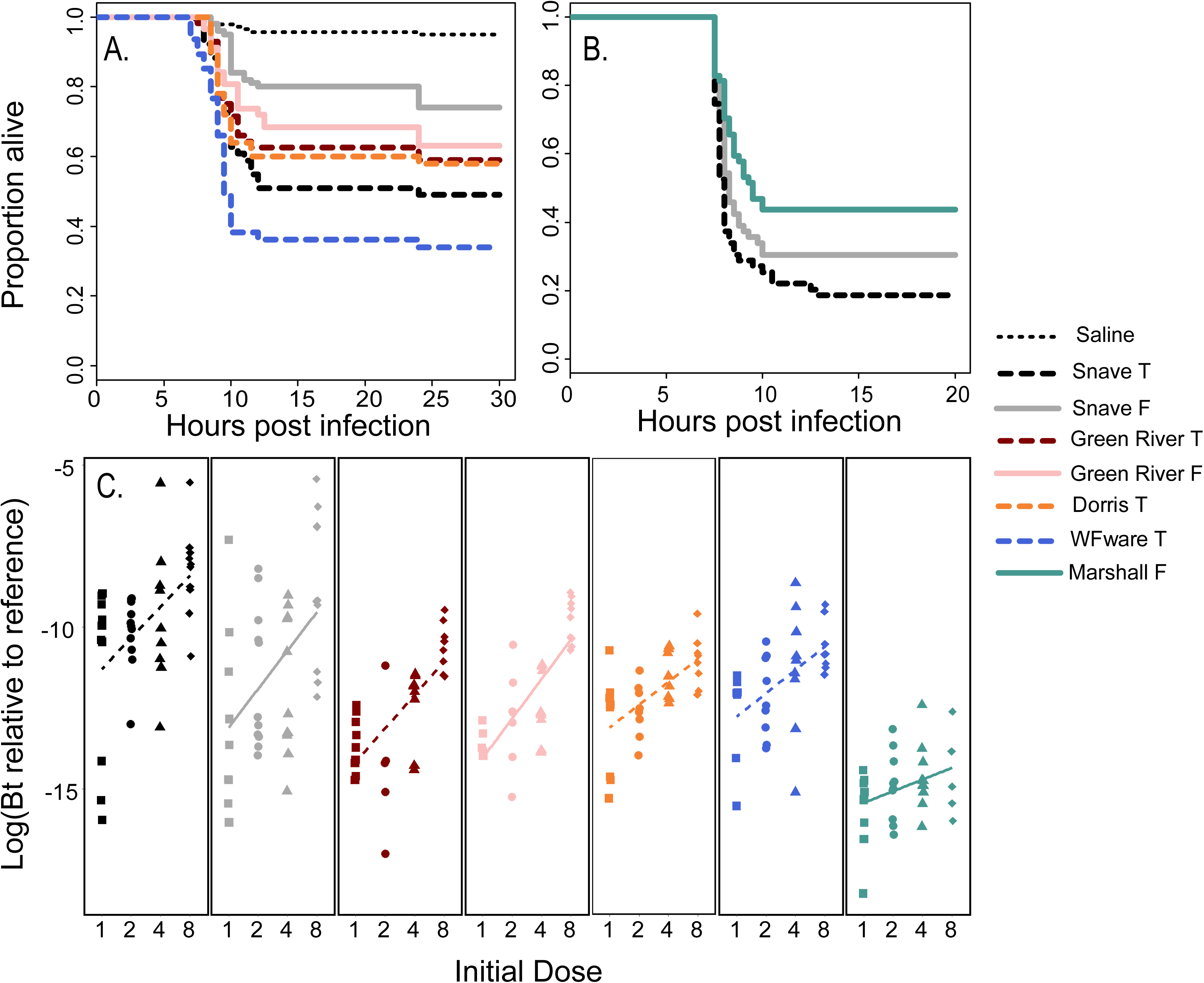
Natural variation in survival and bacterial density during Bt infection. Survival during acute infection with dose “2” was monitored for at least 20 hours post infection in one of two experimental blocks (**A, B**; N = 50-60 beetles/population). To quantify variation in bacterial density, flour beetles were given 2-fold increasing initial doses of Bt (1, 2, 4, 8) and sacrificed 8 hours later. Relative bacterial density for each individual within each population and dose, as quantified by RT-qPCR, is calculated as the log of the linearized difference between Bt-specific and host reference gene expression (**C**). Linear regression lines are provided to visualize increasing bacterial density at 8 hours post infection by initial dose. Populations are color-coded; *T. castaneum* (T) populations have dashed lines and *T. confusum* (F) populations have solid lines.

#### Experiment 3: Survival, bacterial density, and host gene expression after P.lum infection

*Photorhabdus luminescens* is an entomopathogenic symbiont of parasitic nematodes. In nature, it is introduced into the insect hemolymph after the nematode penetrates the cuticle, and is responsible for suppressing host immune responses (Ciche & Ensign 2003) and killing the host so that that nematode can complete its life cycle. To infect beetles with *P. luminescens* (P.lum, ATCC 29999), we prepared an overnight culture from glycerol stocks of the bacterium in Nutrient Broth at 30 °C. We combined 500 uL of log phase culture (OD = 0.405) and 500uL of overnight culture (OD = 1.46), centrifuged it and resuspended the pellet in 450uL of NB, which delivered approximately 250 CFU per needle prick, as determined by plating needle contents. For bacterial and gene expression assays (N = 10 naïve control, N = 10 NB control, N = 20 P.lum infection per population), all beetles were sacrificed at 14 hours post infection by flash freezing and stored at −80°C until processing. For survival assays (N = 50-60/population), individual mortality was monitored every 30 minutes from 12-24 hours post infection (peak mortality 16-20 hours).

#### Experiment 4: The effect of IMD pathway signaling on immune sensitivity

We obtained primer sequences for *T. castaneum imd* dsRNA template from the iBeetle Database (Dönitz *et al*. 2014) (**Table S1**). We produced T7 promoter sequence-tagged DNA via PCR (Platinum Green Hot Start kit, Invitrogen) from *T. castaneum* cDNA, and purified the product using the QIAquick PCR Purification kit (Qiagen), followed by overnight dsRNA synthesis (Megascript T7 kit, Invitrogen) as described in (Posnien *et al*. 2009). We similarly created dsRNA against a maltose binding protein E (malE) sequence using *E. coli* DNA as a template (Yokoi *et al*. 2012a) to serve as a control for the effect of RNAi induction. We quantified knock-down efficiency (87%) by comparing *imd* expression in IMD-RNAi and MalE-RNAi groups relative to the RPS18 reference gene. There was no significant difference in constitutive *imd* expression among MalE-RNAi beetles and completely naïve beetles.

We obtained a cohort of Snavely *T. castaneum* larvae and assigned them to either a control group injected with MalE dsRNA or an *imd-*RNAi group. We injected them with 0.5ug dsRNA dissolved in 1uL sterile insect saline + green dye, as described in (Posnien *et al*. 2009). Three days later, we assigned undamaged individuals from each RNAi treatment group to a sterile saline control or one of four Bt doses as in Experiment 1 (N = ∼ 8 individuals/dose/RNAi treatment), and sacrificed them at 8 hours post injection. We excluded two individuals from the *imd-*RNAi group from the analysis because they had not received proper dsRNA injections, as evidenced by normal *imd* expression. We infected an additional 60 individuals from the MalE and IMD treatment groups with dose 2 (approximately 100 CFU) to monitor survival over 24 hours post Bt infection. We repeated dose1 infections again (50 CFU, N = 11/group) and sacrificed individuals at 6 hours post infection to obtain more normally distributed bacterial density estimates.

### RT-qPCR Primer Design

We chose immune genes to assay based on a previous study that suggests that their expression is correlated to Bt density (Tate & Graham 2017) and where we know the humoral pathways that modulate their expression: the IMD pathway (the AMP *attacin-1* (TC007737) and the recognition protein *pgrp-sc2* (TC013620)), the Toll pathway (the AMP *cecropin-3* (TC000500)), or both (the AMP *defensin-1* (TC006250)) in *T. castaneum* (Koyama *et al*. 2015; Yokoi *et al*. 2012a). Additionally, we assayed *dopa decarboxylase* (*ddc*, TC013480) expression because it is an important enzyme in the melanization pathway (Huang *et al*. 2005), although our results subsequently suggested that its expression may be correlated to IMD pathway-regulated genes (**Fig. S1**) and therefore may not accurately represent melanization dynamics. A primary concern of primer design was to ensure that each primer set amplified the intended target to the same efficiency and Ct range in both *T. castaneum* and *T. confusum*. We designed several degenerate primer sets per gene by comparing CDS sequences from NCBI for *T. castaneum* against *T. confusum* draft sequences provided by Jeffery Demuth (personal comm.). We tested the efficiency of each primer set against cDNA derived from Snavely *T. confusum* and *T. castaneum* individuals. To capture reference gene expression, we used the RPS18 primer set provided in (Lord *et al*. 2010), which had previously demonstrated that the expression of this gene is constant across stages of fungal infection. Primers that performed with high efficiency and equivalently in both species are listed in **Table S1**. We used previously published primers (Tate *et al*. 2017; Tate & Graham 2015b) to quantify Bt density relative to host tissue (**Table S1**), where the amplified product has been previously shown to correlate strongly with bacterial CFU as determined by samples plated on agar (Tate & Graham 2015b).

### Nucleic Acid Processing and RT-qPCR

We extracted total RNA from all individual beetles using the RNeasy Mini Kit (Qiagen) according to the manufacturer’s instructions. We reverse-transcribed 40-80ng RNA to cDNA using quarter-reactions of the Superscript IV VILO MasterMix kit (Invitrogen), and diluted cDNA with 40uL nuclease-free water. Within each experiment, we performed qPCR for each transcript template using PowerUp Sybr Green Master Mix (Life Technologies) on the QuantStudio 6 machine (Applied Biosystems) by running all individuals on the same 384-well plate in duplicate, where possible, using default settings unless indicated in Table S1. We confirmed equivalent Ct thresholds and ran a subset of individuals on both plates to minimize plate effects, as in (Tate *et al*. 2017).

### Statistical Analysis

For the qPCR data, we calculated ΔCt (cycle quantification) values (target gene Ct – reference gene Ct) for each gene template for each individual. These values were linearized (2^−ΔCt^) prior to analysis and then log10-transformed for normality, as in (Tate *et al*. 2017; Tate & Graham 2017). These data are available through the Dryad Digital Repository (Accession XXX). To compare constitutive (naïve individuals) and microbe recognition-independent inducible responses (NB-injected control groups (dose 0)) across populations, we compared gene expression values in the using linear models (“lm” or “aov” functions in R depending on desired output; model = relative expression ∼ treatment or population). Models of immune sensitivity and magnitude, where relative immune gene expression is the dependent variable, took the form of: relative expression ∼ bacterial density + treatment + bacterial density * treatment, where data from all doses were used. For the latter model, we interpreted a significant main effect of treatment as indicating an overall difference in expression independent of microbe density, while a significant microbe density term indicated that the expression of the gene was sensitive to microbe density. A significant interaction term indicated differences in immune sensitivity (the slope of microbe-dependent expression) among treatments or populations. Because we were assaying multiple genes, we applied a Bonferroni correction of (α = 0.05/N), where N is the number of assayed genes, as specified in the table notes.

To compare survival among populations, we employed censored Cox Proportional Hazards tests to estimate variation among populations and to obtain hazard ratios for all populations relative to Snavely *T. confusum* (or MalE, for the RNAi experiment). Pairwise correlations among infection mortality hazard ratios and immune parameters were quantified using Pearson correlations (rcorr function in R) and adjusted raw p-values for false discovery rate using the Benjamini-Hochberg method (Benjamini & Yekutieli 2001). However, it should be noted that since we only had seven populations and since many of our immune parameters are co-correlated and independently support similar associations, FDR correction probably inflates Type II error in an overly conservative manner for any given pairwise test.

## Results

### Host survival and resistance to bacterial infection vary among populations and species

Survival rates during the acute phase of septic Bt infection differed significantly among beetle populations (**Fig. 1A, B**). We calculated hazard ratios for each population relative to the Snavely *T. confusum* population as it provided maximal differentiation among populations (censored Cox Proportional Hazards, survival ∼ site + experimental block, N = 50-60 beetles/population). The other two *T. confusum* populations did not differ significantly from Snavely *T. confusum* (Marshall HR = 0.67, p = 0.084, Green River Hazard Ratio = 1.51, p = 0.16), but all four *T. castaneum* populations were significantly more likely to die (Green River HR = 1.83, p = 0.035; Dorris HR = 2.01, p = 0.017; Snavely HR = 2.53, p < 0.001; WF Ware HR = 4.21, p < 0.0001). Sterile saline injections resulted in negligible mortality (7/140 beetles, or 5%; **Fig. 1A**), which was evenly distributed among different populations.

Beetle populations also varied in their resistance to Bt infection, as measured by RT-qPCR at 8 hours post infection with one of four 2-fold increasing doses (N = 6-10 beetles/dose/population). Relative Bt density at 8 hours post infection differed by 4-5 orders of magnitude for any given initial dose between the most resistant (Marshall *T. confusum*) and least resistant (Snavely *T. castaneum*) populations (**Fig. 1C**). There was a strong correlation (R^2 = 0.91, p = 0.0045) between lowest and highest initial doses for average 8 hour bacterial density among populations. Overall, *T. confusum* populations were significantly more resistant to Bt than *T. castaneum* populations (Linear model, log(bacterial density) ∼ species + dose, estimate = −1.24, st. error = 1.83, t = 4.7, p < 0.0001), and bacterial load at 8 hours post infection increased significantly with initial dose (est. = 0.964, st. error = 0.12, t = 8.13, p < 0.0001). Population-level bacterial density averages at 8 hours post infection (from both low and high initial doses) were correlated to population survival hazard ratios (dose 1: R^2^ = 0.72, p = 0.071; dose 4: R^2^ = 0.73, p = 0.061), although neither pair-wise correlation was significant.

Survival rates against *P. luminescens* were more homogenous among populations (**Fig. S2**), although Marshall *T. confusum* was once again the least susceptible (50% survival at 24 hours post infection; N = 50-60 beetles/population). Relative to Marshall, only Green River *T. confusum* was significantly less likely to survive (HR = 1.93, p = 0.010), although WF Ware *T. castaneum* also did fairly poorly (HR = 1.58, p = 0.081). Once again, sterile saline injections resulted in negligible mortality (2/55 beetles, or 3.6%).

During the first 14 hours after infection, *P. luminscens* bacterial density increased by an average of 4.5 orders of magnitude, but bacterial density variation among populations (**Fig. S2**) was more homogenous than observed with Bt. Marshall *T. confusum* was once again among the most resistant populations, but only Snavely *T. castaneum* had a significantly higher bacterial load than Marshall (estimate = 10^2.26, t = 2.3, p = 0.0284). Average bacterial density at 14 hours post infection was completely uncorrelated to mortality hazard ratios (R^2 = 0.01, **Fig. S3**).

### Inducible immunity is sensitive to microbe density but sensitivity varies by population

To examine variation in constitutive immunity among populations (**Fig. 2A, B**), we quantified the magnitude of gene expression in individuals that were stabbed with saline but then immediately sacrificed (within 2 min). All assayed genes exhibited significant variation among populations (ANOVA, *attacin-1*: F_6,80_ = 3.22, p = 0.0069; *defensin-1:* F_6,80_ = 21.05, p < 0.0001; *pgrp-sc2*: F_6,80_ = 8.155, p < 0.0001). Populations fell roughly into low (significantly different from Snavely *T. castaneum*) and high expression bins, where the former includes Snavely *T. confusum*, Dorris *T. castaneum*, WF Ware *T. castaneum*, and Marshall *T. confusum*, while the higher expression group includes both *T. castaneum* and *T. confusum* from Green River (**Table S2, Fig. 2B**).

**Figure 2.**
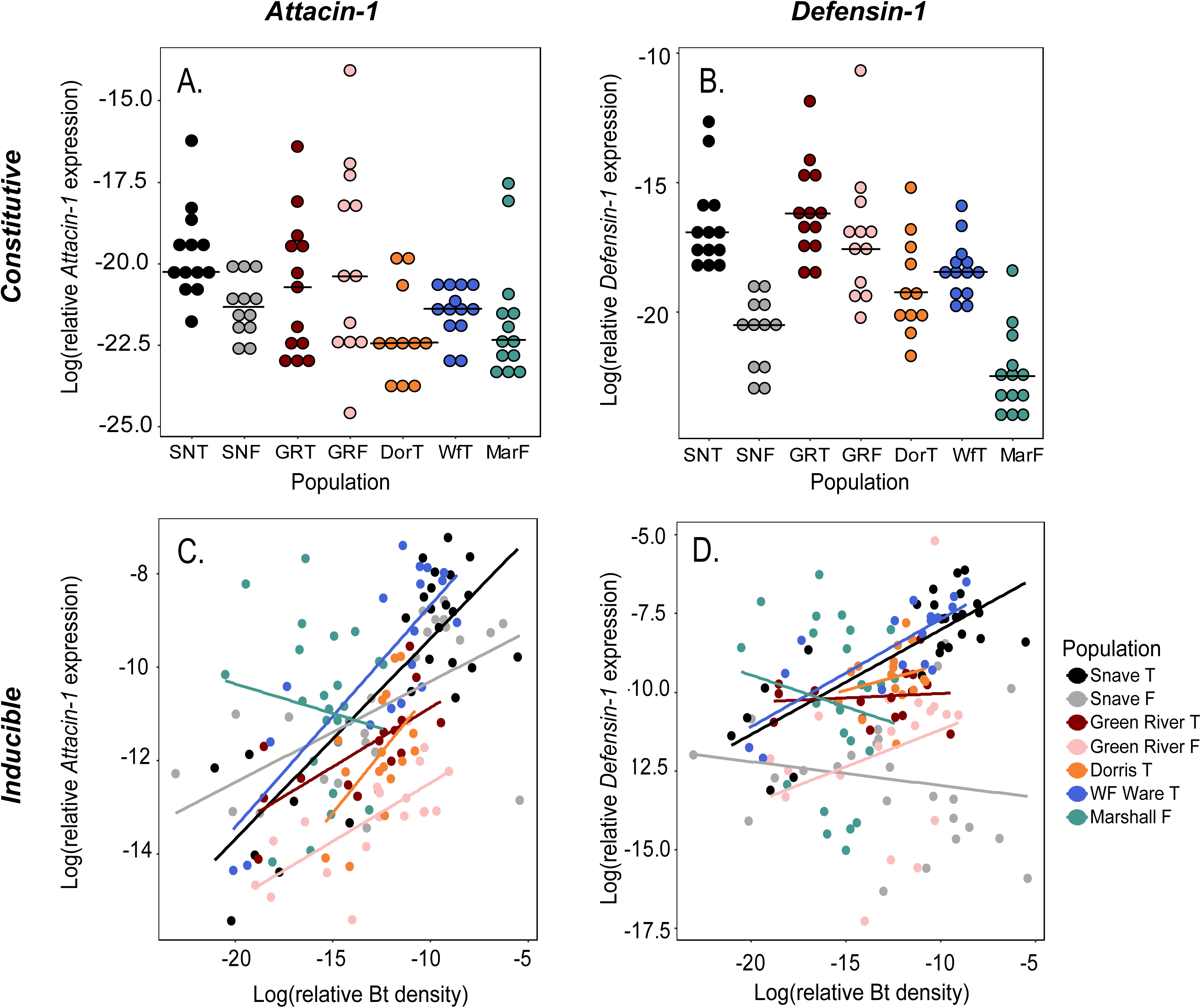
Patterns of constitutive and inducible *attacin-1* and *defensin-1* expression by population. As quantified by RT-qPCR, the log of relative immune gene expression prior to infection varied among naïve individuals from each population (*attacin-1*, **A**.; *defensin-1*, **B**.) Black lines indicate median values for each population. The relationship between the log of bacterial density and the log of *attacin*-1 (**C**.) and *defensin-1* (**D**.) gene expression 8 hours after challenge with saline or Bt infection illustrates variation in both the intercept (microbe-independent) and slope (microbe-dependent sensitivity) of inducible immune gene expression among populations. Lines represent linear fits for each main variable level as computed by the “lm” function in the geom_smooth algorithm of ggplot2 (R).

To quantify the magnitude of microbe-independent investment in inducible immunity, we compared the expression of immune genes 8 hours after induction with a sterile saline jab. For both *attacin-1* and *defensin-1*, the main effect of population was significant (ANOVA, *att-*1 F_6,26_ = 3.31, p = 0.015; *def-1* F_6,26_ = 3.655, p = 0.0091). Among most populations the range of both *attacin-1* and *defensin-1* expression spanned around 2.5 orders of magnitude, but Marshall exhibited significantly higher expression of both genes than the other populations (**Table S2**).

To quantify microbe density-dependent responses at 8 hours post challenge (combined saline and Bt-infected; N = 20-25 individuals/population, **Fig. 2C, D**), we employed a linear model of the form: expression ∼ population + bacterial density + population*bacterial density. Bt density strongly predicted expression of all assayed genes at 8 hours post infection (**Table 1**). The wild-derived populations varied significantly in microbe-sensitive expression (Bt density by population interaction) of *attacin-1* (**Fig. 2C**), *defensin-1* (**Fig. 2D**), and *pgrp-sc2* (**Table 1, Fig. S4**), but not *ddc* (**Fig. S4**; F_6,129_ = 1.383, p = 0.226). The Marshall *T. confusum* population stood out as having qualitatively different expression patterns relative to other populations (**Fig. 2**). The other *T. confusum* populations conformed to general sensitivity patterns, except that they demonstrated a bimodal distribution of *defensin-1* expression where some expressed it at high levels at any given microbe density and some expressed it at much lower levels (**Fig. 2**).

**Table 1.**
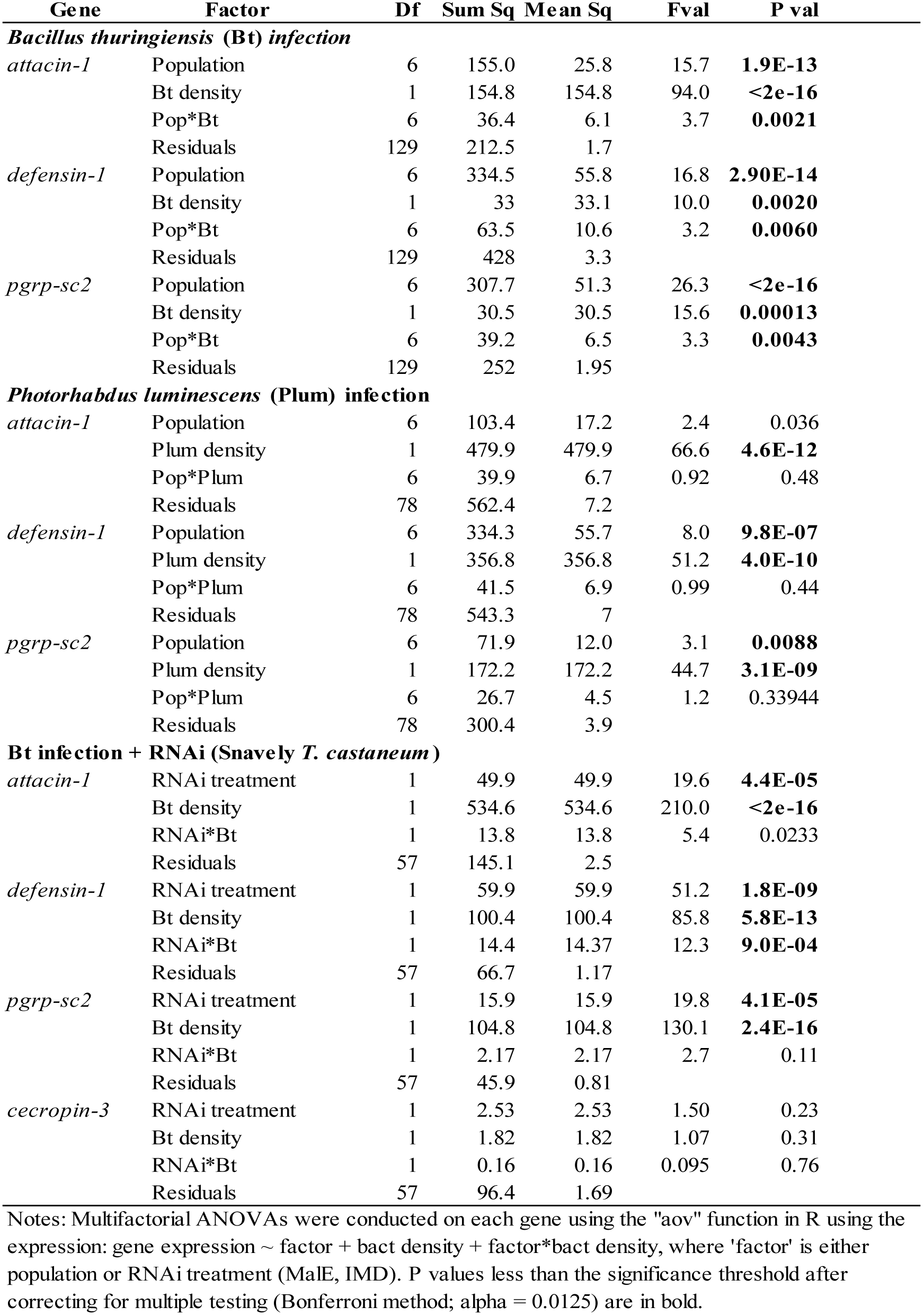
Immune gene expression sensitivity to bacterial density, population, and their interaction.

All assayed genes showed significant variation in expression among populations 14 hours after *P. luminescens* infection (**Table 1, Fig. S2**, ANOVA, N = 14-16 individuals/population), and the expression of all genes was significantly correlated to bacterial density (**Table 1**). However, there was no significant variation in the sensitivity of gene expression to bacterial density among populations (**Table 1**).

### Bacterial load and gene expression at time of death varies to some extent by population

As the bacterial load at time of death (BLUD) has been previously proposed as a measure of infection tolerance in fruit flies (Duneau *et al*. 2017), we sacrificed a subset of beetles from the Bt infection experiment as soon as they exhibited moribund behavior to quantify bacterial load and gene expression. Populations showed modest but significant variation in bacterial load at time of death (**Fig. S5A**), which was not predicted by post-infection time of death (**Table S4, Fig. S5B**). There was substantial variation in bacterial load at time of death within populations (**Fig. S5A**). Populations differed significantly in immune gene expression at the time of death (**Table S4**). For example, all *T. confusum* populations relative to *T. castaneum* had lower expression levels of *defensin-1* at death (**Fig. S5C**), and WF Ware *T. castaneum* exhibited lower expression of the melanization enzyme *ddc* (**Fig. S5D**). Immune gene expression was still significantly correlated to Bt density (**Table S4**), but the slope was gently negative (**Fig. S5C, D**), and the population-by-Bt density interaction terms were not significant for any of the genes (**Table S4**).

### Microbe-independent and -dependent inducible immune parameters are associated with host survival

To uncover associations between infection survival, bacterial resistance, and immune parameters, we calculated population averages for microbe-independent constitutive and inducible gene expression as well as inducible sensitivity (the slope of gene expression by Bt density) for *attacin-1, defensin-1*, and *pgrp-sc2*. We performed pairwise Pearson correlation analysis on these parameters against Bt and *P*.*lum* survival hazard ratios, Bt density, *P*.*lum* density, and Bt density at time of death (**Fig. 3A**; circle presence indicates R^2^ > 0.75 and p < 0.05 before FDR correction; none but Bt dose 1 × Bt dose 4 were significant after FDR correction). While microbe-independent inducible *defensin-1* (and *attacin-1*) magnitude did not correlate with Bt-induced mortality (R^2^ = 0.05, **Fig. 1B**), it did show a negative correlation with P.lum-induced mortality (R^2^ = 0.84, **Fig. 3C**). On the other hand, the steepness of the microbe-dependent slope of *defensin-1* induction was positively correlated with Bt-induced mortality (R^2^ = 0.72, **Fig. 3D**) but not with P.lum-induced mortality (R^2^ = 0.54, **Fig. 3E**). The bacterial load at time of death (BLUD) was not associated with any of the immune parameters, and did not correlate to mortality rates. All pairwise correlation R^2^ values are available in **Fig. S3**.

**Figure 3.**
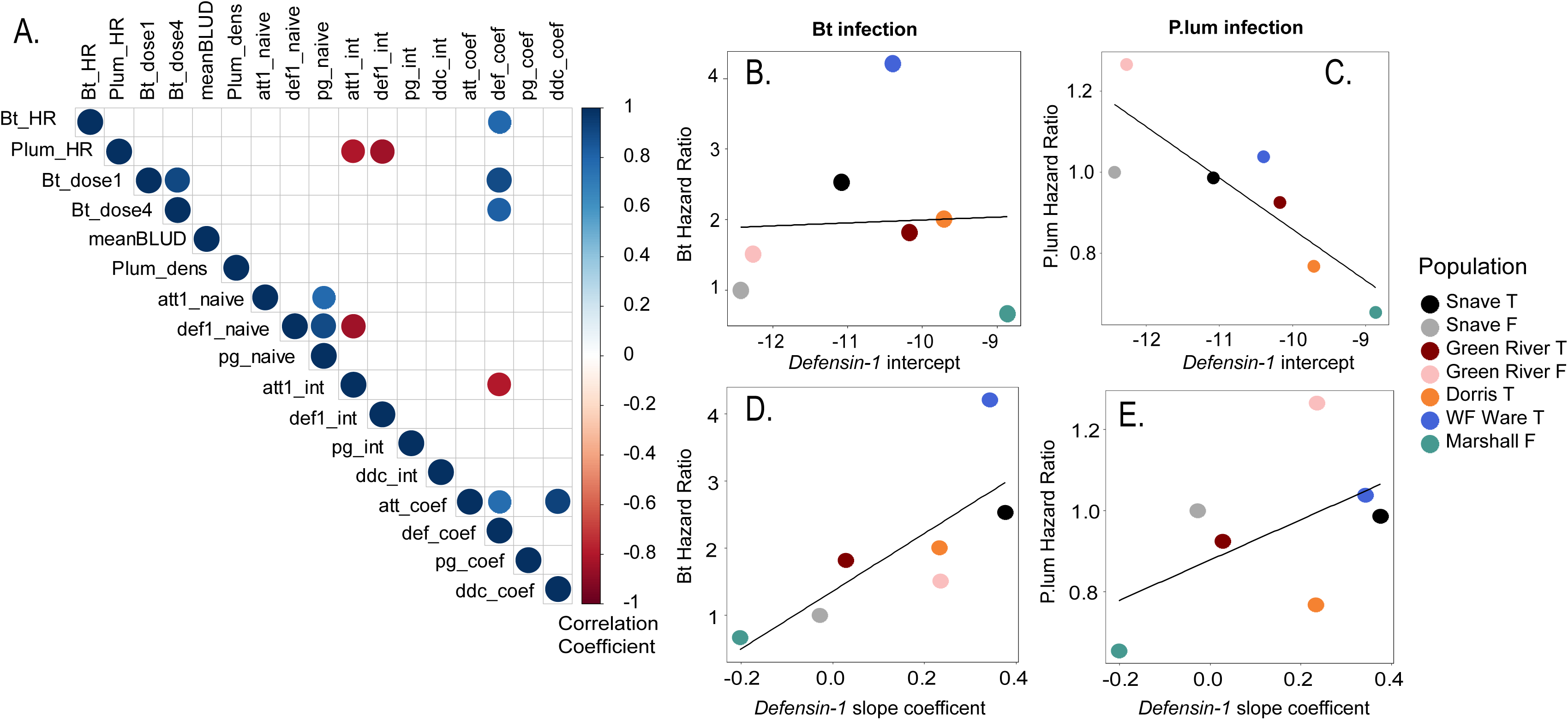
Correlations between constitutive and inducible immune parameters and phenotypic outcomes of infection with *Bacillus thuringiensis* and *Photorhabdus luminescens*. Quantification of Pearson correlation coefficients (significant pairwise correlations, **A**) identified no relationship between the microbe-independent inducible intercept of *defensin-1* expression and Bt mortality (**B**, as quantified by population Hazard Ratios), but a significant negative relationship between *defensin-1* intercept and mortality from *P. luminescens* infection (**C**). On the other hand, the microbe-dependent slope coefficient of *defensin-1* expression was significantly correlated to Bt-induced mortality (**D**), with a positive but non-significant association with *P. luminescens* mortality (**E)** as well. HR = hazard ratio, BLUD = Bt density at death, Bt_dose and Plum_dens = bacterial density at 8 and 14 hours post infection respectively, naïve = expression in uninfected individuals, int = microbe-independent inducible gene expression intercept (saline-injected expression at 8 hours), coef = slope of expression over bacterial density. Att1 = *attacin-1*, Def1 = *defensin-1*, pg = *pgrp-sc2*, ddc = *dopa decarboxylase*. Lines represent linear fits for each main variable level as computed by the “lm” function in the geom_smooth algorithm of ggplot2 (R).

### Knock-down of imd signaling affects bacterial resistance and inducible immune sensitivity

In the full dose-response experiment (**Fig. 4**), RNAi-mediated knock-down of *imd* expression (4-fold reduction relative to MalE-injected individuals) in Snavely *T. castaneum* did not significantly affect bacterial density in infected individuals even as dose was a highly significant predictor of Bt density (linear model, IMD vs. MalE, effect of treatment: estimate = 10.7, st. error = 0.78, t = 1.322, p = 0.191; effect of dose: estimate = 2.44, st. error = 0.26, t = 9.19, p < 0.00001). However, we noticed that bacterial density followed a non-continuous distribution of low and high groups within and among doses (**Fig. 4C)** consistent with bifurcating infection outcomes ((Duneau *et al*. 2017) and complicating statistical interpretation. Therefore, we performed another infection experiment with a lower dose and earlier sampling to avoid the bimodal distribution (**Fig. 4B**), and in this experiment IMD-RNAi individuals had a significantly higher bacterial load than MalE-RNAi individuals (linear model, estimate = 18-fold increase in Bt density, st. error = 1.88, t = 4.556, p = 0.000378). Despite the exacerbation of bacterial density in IMD-RNAi individuals, this group was not more likely to die during the acute infection phase relative to MalE-RNAi individuals (**Fig. 4A**, Cox Proportional Hazards, N = 50/treatment, hazard ratio = 1.05, Z = 0.174, p = 0.86).

**Figure 4.**
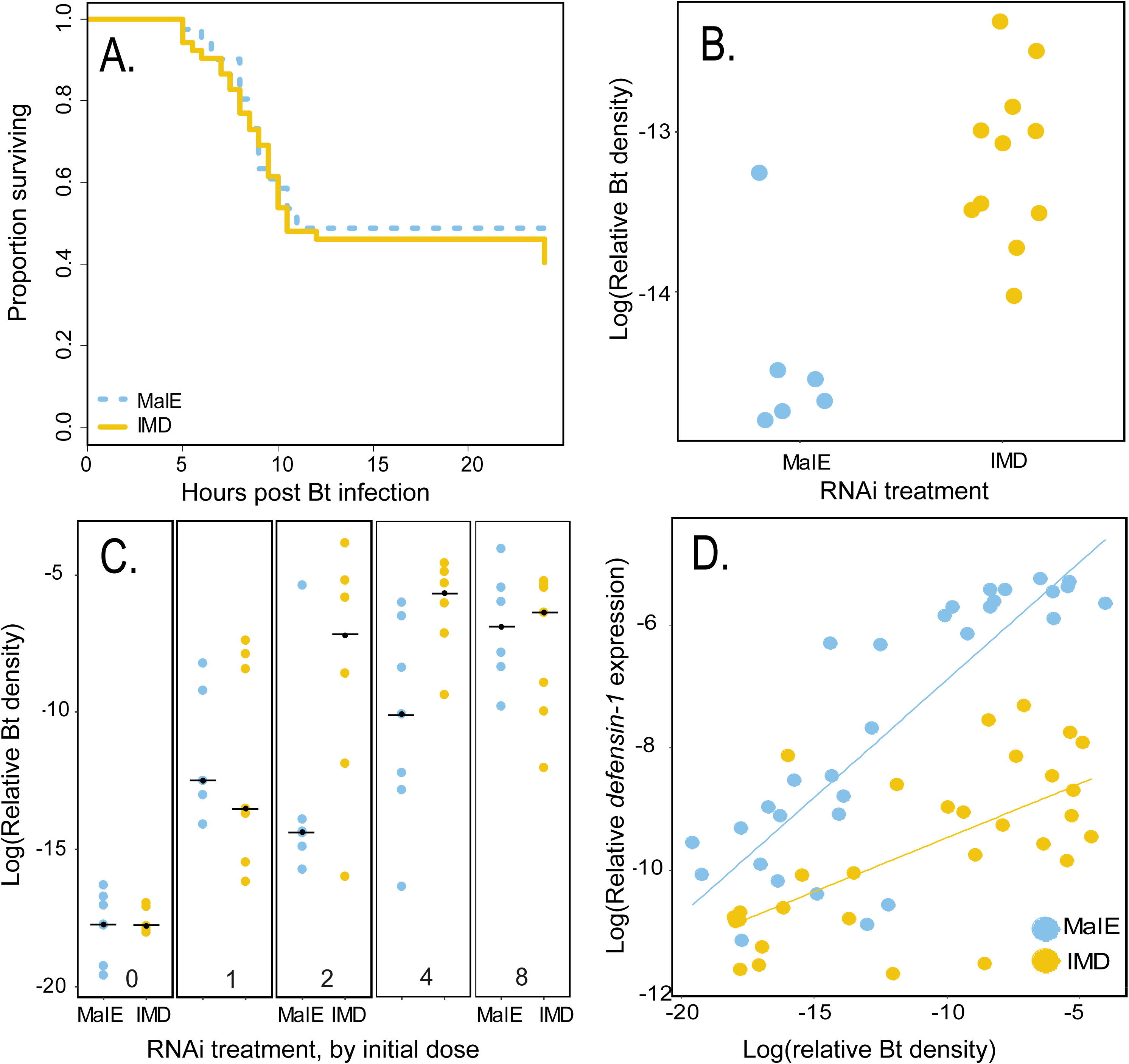
RNAi-mediated knockdown of *imd*, a regulatory element of the IMD pathway, impacts bacterial density and inducible immune parameters but not survival against Bt infection. Snavely *T. castaneum* individuals were injected with dsRNA against the *imd* gene (IMD, yellow), or a no-target dsRNA injection control (MalE, blue). Individuals were injected with an LD50 dose of Bt and the proportion surviving was monitored for 24 hours (**A**). Individuals were administered a low dose (**B**) or increasing doses (**C**) of Bt, and relative bacterial density was quantified via RT-qPCR at 8 hours post infection. Black dots represent the median bacterial density for each group. These same individuals (as well as saline controls) were also assayed for expression of *defensin-1* relative to bacterial density (**D**) to quantify the impact of the imd pathway on microbe independent and dependent inducible immune responses. Lines represent linear fits for each main variable level as computed by the “lm” function in the geom_smooth algorithm of ggplot2 (R).

In the full dose-response experiment (**Fig. 4D**; ANOVAs in **Table 1**), the knockdown of *imd* significantly reduced the mean magnitude of *attacin-1* and *defensin-1* expression after Bt infection. IMD knock-down also significantly reduced the sensitivity of *defensin-1* expression to Bt density (RNAi treatment × Bt density interaction) without negating the main effect of Bt density on gene expression.

## Discussion

The parasites to which hosts are exposed, the frequency of exposure, and the costs and benefits associated with mounting defenses against those parasites, are all expected to contribute to the evolution of inducible immune responses (Cressler *et al*. 2015; Frank 2002; Hamilton *et al*. 2008; Mayer *et al*. 2016). The dynamics of immunity during the acute infection phase are particularly important for host fitness, as modest levels of variation in early responses could lead to drastic variation in host mortality and parasite persistence (Duneau *et al*. 2017). However, the ecological, microbial, and temporal factors to which these dynamics are sensitive have not been well-established, providing few avenues to quantify and understand natural variation in inducible immune responses. Recent experiments have suggested that immune effector expression in insects might indeed be sensitive to microbe density (Louie *et al*. 2016; Tate & Graham 2017), raising a host of new questions surrounding the dynamics of inducible immune responses. In this study, we discovered substantial variation in both microbe-dependent and independent inducible immune dynamics among flour beetle populations during acute infection with Bt, suggesting evolutionary maintenance of variation in both traits. The average magnitude of microbe-independent inducible immunity at the population level was positively associated with survival against *P. luminescens*, a slow-growing but virulent bacterium that inhibits host inducible responses (Hwang *et al*. 2013). On the other hand, the slope of microbe-sensitive inducible *defensin-1* expression was associated with poor survival outcomes against Bt. As expected, interfering with inducible immune signaling via RNAi-mediated knockdown of *imd* reduced the magnitude of microbe-independent inducible immune gene expression, but for *defensin-1* it also affected microbe-dependent sensitivity. This study provides the first evidence, to our knowledge, of natural variation in these facets of inducible immune dynamics and suggests that these dynamics may be associated with different costs and benefits when confronted with microbes that vary in antigenic and pathogenic life history traits.

### The costs and benefits of immunological sensitivity to microbe density

Literature from as far back as the 1950s testifies to the diversity of bacteria, (Burges & Weiser 1973), protozoa (Park & Marian Burton 1950), microsporidia (Milner 1972), fungi (Burges & Weiser 1973), and helminths (Yan *et al*. 1998) with which flour beetles are naturally infected. Given that different populations probably face evolutionary pressures from different subsets of these parasites, it is not surprising that our seven populations demonstrated substantial natural variation in immune system activation, resistance, and survival outcomes against two focal microbes.

From an evolutionary perspective, should we expect greater variation in microbe-independent responses than in microbe-sensitive ones? An on/off binary response to microbial exposure should be associated with the same kinds of energetic and immunopathological sunk costs that we traditionally associate with constitutive immunity, while a response that is able to respond dynamically to changes in microbe density would allow greater flexibility to pay costs only when needed, as well as maintain populations of commensals or beneficial microbes. However, the latter strategy risks losing control over fast-growing microbes. Optimal investment in microbe-independent and sensitive inducible dynamics is also likely to depend on the frequency and predictability of parasite exposure, complementing predictions made for optimal investment in constitutive and inducible immunity (Hamilton *et al*. 2008; Mayer *et al*. 2016). Finally, epidemiological feedbacks in populations with different immune strategy distributions should lead to different pressures on transmission and virulence (Cressler *et al*. 2015), further creating an opportunity for natural variation in both microbe dependent and independent terms.

In this study, we focused on two AMPs that are sensitive to microbe density in *T. castaneum* (Tate & Graham 2017). Previous work has established that the AMP *attacin-1* is regulated by the IMD pathway in *T. castaneum* (Yokoi *et al*. 2012a). The AMP *defensin-1* is also regulated by IMD signaling, but is additionally co-regulated by the Toll pathway and other tissue-dependent mechanisms (Tzou *et al*. 2000; Yokoi *et al*. 2012b). Thus, while both are sensitive to microbe density and thus correlated (**Fig. S1**; (Tate & Graham 2017)), we did not expect them to be entirely co-regulated. In our study, two *T. confusum* populations (Snavely and Marshall) that showed heightened survival against Bt relative to other populations (**Fig. 1A, B**) also showed lower *defensin-1* sensitivity to microbe density (**Fig. 2D**), echoing a broader association between mortality and *defensin-1* sensitivity among our populations (**Fig. 3D**).

This does not imply that *defensin-1* is somehow intensely immunopathological and contributing causally to mortality. Instead, we suspect that *defensin-1* sensitivity may simply covary with a suite of stress responses that indicate failure to control infections, including high damage signals, oxidative stress, and general pathology. Supporting this idea, knocking down *imd* reduces *defensin-1* expression by around three orders of magnitude at the highest bacterial densities (**Fig. 4D**), reduces sensitivity, and reduces resistance by at least one order of magnitude (**Fig. 4B, C**) but does not ultimately impact survival or even the distribution of time to death in insects inoculated with an LD50 dose of Bt (**Fig. 4A**). It may be that IMD pathway-mediated immunopathology is equivalent to approximately one order of magnitude of bacterial virulence per unit time, thus producing no net effect on host survival. It is more likely, however, that survival against Bt is determined long before the inducible immune response becomes relevant, relying instead on variation in alternative arms of the immune system like melanization or phagocytosis to create differences in Bt-induced mortality rates among individuals and populations (Duneau *et al*. 2017). Finally, AMP expression was sensitive to *P. luminescens* density, but there was very little variation in sensitivity among populations just as there was negligible variation in resistance and mortality rates (**Fig. S1**). Thus, we propose that the microbe-dependent inducible immune gene expression parameter may be a quantifiable marker of overall infection pathology and virulence.

On the other hand, the microbe-independent inducible immune parameter may be functionally relevant against slower-growing but manipulative microbes, as the average expression of both *attacin-1* and *defensin-1* at 8 hours post saline-stab was associated with slower mortality rates during *P. luminescens* infection (**Fig. 3A, C**). This is not particularly surprising, since the bacterium goes through the trouble of suppressing AMPs in its native lepidopteran hosts (Nielsen-LeRoux *et al*. 2012) and thus AMPs must present at least some threat to the bacterium. Earlier and stronger expression of AMPs before bacteria reach high densities could help delay bacterial growth and the onset of density-dependent virulence factor production through quorum-sensing, a hallmark of *Photorhabdus* and Bt life history strategies (Nielsen-LeRoux *et al*. 2012).

### Bacterial load at time of death and its association with infection tolerance

The bacterial load at time of death (BLUD) has been proposed as a metric of infection tolerance (Duneau *et al*. 2017) since it should take more bacteria to kill a more tolerant host, all other things being equal. For example, the average BLUD of *D. melanogaster* adults is lower upon infection with more virulent bacteria relative to less virulent bacteria, and is independent of the time to death (Duneau *et al*. 2017). In our study, the BLUD of larval *Tribolium* beetles was also uncorrelated to time of death after infection with Bt (**Fig. S5B**). However, there was substantially more variation in BLUD among individuals within populations than there was among populations (**Fig. S5A**), and average BLUD was not associated with any survival metrics at the population level (**Fig. S2A**), clouding its utility as a proxy of infection tolerance. Whether variation in BLUD reflects variation in factors associated with tolerance, such as damage repair, energetic provisioning, or immunopathology (Ayres & Schneider 2012), still requires further study in this and other species.

It is worth noting that AMP expression sensitivity to bacterial load in moribund individuals was dampened or even slightly negative (**Fig. S5C**), and while sensitivity did not significantly differ among populations, the overall magnitude of *defensin-1* expression at time of death was lower in *T. confusum* populations relative to *T. castaneum* (**Fig. S5C**). Louie *et al*. (Louie *et al*. 2016) found that different AMPs show different sensitivities to *Listeria monocytogenes* density in fruit flies recovering with the aid of antibiotics, and that unlike our acute infection phase data, the sensitivity relationship had a decidedly sigmoidal shape that included a maximal expression level. Collected well prior to the onset of host mortality for both bacterial species, our acute phase data likely reflect the linear portion of the sensitivity curve, but at the time of death individuals may have hit the flat maximum of the sigmoid. Our data suggest that the intercept and linear slope of this curve are likely to be relevant for infection outcomes; whether the maximum expression level has functional consequences for host survival or microbial transmission represents an avenue for future study.

### Future directions

Our observation that *T. confusum* populations are more likely to survive Bt infection than *T. castaneum* populations echoes a classic observation of heightened resistance to coccidian infection in *T. confusum* relative to *T. castaneum* that modulated competitive dynamics among the two host species (Park & Marian Burton 1950). As these species co-occur in many temperate regions, including two sites (Green River and Snavely) that we sampled for this study, it would be interesting to determine the extent to which variation in microbial sensitivity arises through local adaptation as opposed to species-level variation in immune system architecture, and to what extent inducible immune variation could influence competition and coexistence among host species in communities that share parasites.

The mechanisms that control microbe density-dependent sensitivity remain unclear. Knock-down of *imd* expression using RNAi reduced the mean expression of *attacin-1* and *defensin-1* during Bt infection by over three orders of magnitude but only affected the sensitivity of *defensin-1* (**Table 1**). It is possible that targeting recognition protein abundance or the affinity of transcription factors for AMP-specific promoter regions might produce variation in the sensitivity of AMP expression to microbe density, and thus represent avenues for future manipulation. It is also possible that sensitivity is driven by spatial considerations, for example the rate of bacterial dissemination throughout the hemocoel, but it is not clear why this would differ among species and populations. Uncovering the mechanism(s) regulating inducible sensitivity to microbe density, coupled with experimental evolution of sensitivity, would allow the estimation of the relative costs and benefits of microbe density-dependent and independent inducible immune responses.

### Conclusions

In this study, we have demonstrated that flour beetle populations exhibit variation in both microbe density-dependent and –independent inducible immune parameters, and that the relative costs and benefits of investment in inducible immunity are relevant for understanding infection outcomes in a world where hosts are assailed by a diversity of unpredictable parasites. We recommend that theoretical studies on immune system evolution or host-microbe coevolution consider these results when building models and parameterizing the associated fitness costs of immunity. In a similar vein, eco-immunologists should consider multiple immunological parameters when designing experiments to disentangle relationships between infection, the magnitude of immunological investment, and host fitness.

## Acknowledgements

We would like to thank Jeff Demuth for providing *T. confusum* gene sequences and Allison Leich-Hilbun for comments on the draft.

## Data Accessibility

The RT-qPCR data and metadata will be deposited into Data Dryad upon manuscript acceptance.

## Author Contributions

A.T.T. conceived the study, A.T.T., D.G., A.P, J.C. designed the experiments, D.G., A.P., J.C, A.T.T. performed the experiments, A.T.T., A.P., J.C., D.G. analyzed the data, and A.T.T. wrote the manuscript with input from all authors.

## Supplemental Tables (Separate File)

**Table S1:** Primer sequences used in study

**Table S2**: Gene expression in naïve and saline-stabbed beetle populations

**Table S3:** Population hazard ratios and gene expression means for naïve, saline-injected, and bacteria-infected individuals used in correlation plots

**Table S4**: Bacterial load at time of death, correlations with immune gene expression, and population variation

## Supplemental Figures

**Figure S1**. Correlations in the relative expression levels of immune genes and bacterial density for host species, by site.

**Figure S2**. Survival, resistance, and immune gene expression of natural populations during *Photorhabdus luminescens* infection.

**Figure S3**. Correlations between constitutive and inducible immune parameters and phenotypic outcomes of infection with *Bacillus thuringiensis* and *Photorhabdus luminescens*.

**Figure S4**. Patterns of constitutive and inducible immune gene expression.

**Figure S5**. Bacterial density and immune gene expression at the time of infection-induced mortality with Bt.

## References

Ardia DR, Gantz JE, Brent C, Schneider, Strebel S (2012) Costs of immunity in insects: an induced immune response increases metabolic rate and decreases antimicrobial activity. Functional Ecology 26, 732–739.

Ayres JS, Schneider DS (2012) Tolerance of Infections. Annual Review of Immunology 30, 271–294.

Bajgar A, Kucerova K, Jonatova L, et al. (2015) Extracellular Adenosine Mediates a Systemic Metabolic Switch during Immune Response. PLoS Biol 13, e1002135.

Benjamini Y, Yekutieli D (2001) The Control of the False Discovery Rate in Multiple Testing under Dependency. The Annals of Statistics 29, 1165–1188.

Burges HD, Weiser J (1973) Occurrence of pathogens of the flour beetle, Tribolium castaneum. Journal of Invertebrate Pathology 22, 464–466.

Ciche TA, Ensign JC (2003) For the insect pathogen Photorhabdus luminescens, which end of a nematode is out? Appl. Environ. Microbiol. 69, 1890–1897.

Cressler CE, Graham AL, Day T (2015) Evolution of hosts paying manifold costs of defence. Proceedings of the Royal Society of London B: Biological Sciences 282.

Dejnirattisai W, Jumnainsong A, Onsirisakul N, et al. (2010) Cross-Reacting Antibodies Enhance Dengue Virus Infection in Humans. Science 328, 745–748.

Dönitz J, Schmitt-Engel C, Grossmann D, et al. (2014) iBeetle-Base: a database for RNAi phenotypes in the red flour beetle Tribolium castaneum. Nucleic Acids Research, gku1054.

Duneau D, Ferdy J-B, Revah J, et al. (2017) Stochastic variation in the initial phase of bacterial infection predicts the probability of survival in D. melanogaster. Elife 6, e28298.

Frank SA (2002) Immune Response to Parasitic Attack: Evolution of a Pulsed Character. Journal of Theoretical Biology 219, 281–290.

Gupta V, Vale PF (2017) Nonlinear disease tolerance curves reveal distinct components of host responses to viral infection. Royal Society open science 4, 170342.

Hamilton R, Siva-Jothy M, Boots M (2008) Two arms are better than one: parasite variation leads to combined inducible and constitutive innate immune responses. Proceedings of the Royal Society B: Biological Sciences 275, 937–945.

Huang CY, Chou SY, Bartholomay LC, Christensen BM, Chen CC (2005) The use of gene silencing to study the role of dopa decarboxylase in mosquito melanization reactions. Insect Molecular Biology 14, 237–244.

Hwang J, Park Y, Kim Y, Hwang J, Lee D (2013) An entomopathogenic bacterium, Xenorhabdus nematophila, suppresses expression of antimicrobial peptides controlled by Toll and IMD pathways by blocking eicosanoid biosynthesis. Archives of Insect Biochemistry and Physiology 83, 151–169.

Koyama H, Kato D, Minakuchi C, et al. (2015) Peptidoglycan recognition protein genes and their roles in the innate immune pathways of the red flour beetle, Tribolium castaneum. Journal of Invertebrate Pathology 132, 86–100.

Login FH, Balmand S, Vallier A, et al. (2011) Antimicrobial Peptides Keep Insect Endosymbionts Under Control. Science 334, 362–365.

Lord JC, Hartzer K, Toutges M, Oppert B (2010) Evaluation of quantitative PCR reference genes for gene expression studies in Tribolium castaneum after fungal challenge. Journal of Microbiological Methods 80, 219–221.

Louie A, Song KH, Hotson A, Thomas Tate A, Schneider DS (2016) How Many Parameters Does It Take to Describe Disease Tolerance? PLoS Biol 14, e1002435.

Mayer A, Mora T, Rivoire O, Walczak AM (2016) Diversity of immune strategies explained by adaptation to pathogen statistics. Proceedings of the National Academy of Sciences 113, 8630–8635.

Milner RJ (1972) Nosema whitei, a microsporidan pathogen of some species of Tribolium. Journal of Invertebrate Pathology 19, 231–238.

Murphy L, Pathak AK, Cattadori IM (2013) A co-infection with two gastrointestinal nematodes alters host immune responses and only partially parasite dynamics. Parasite Immunology 35, 421–432.

Nielsen-LeRoux C, Gaudriault S, Ramarao N, Lereclus D, Givaudan A (2012) How the insect pathogen bacteria Bacillus thuringiensis and Xenorhabdus/Photorhabdus occupy their hosts. Current Opinion in Microbiology 15, 220–231.

Park T, Marian Burton F (1950) The population history of *Tribolium* free of sporozoan infection. Journal of Animal Ecology 19, 95–105.

Jent D., Perry A., Critchlow J., Tate, A.T. Data from: Natural variation in the contribution of microbial density to inducible immune dynamics. Dryad Data Repository DOI: XXXX

Posnien N, Schinko J, Grossmann D, et al. (2009) RNAi in the Red Flour Beetle (Tribolium). Cold Spring Harb Protoc 2009, pdb.prot5256-.

Raymond B, Johnston PR, Nielsen-LeRoux C, Lereclus D, Crickmore N (2010) Bacillus thuringiensis: an impotent pathogen? Trends in Microbiology 18, 189–194.

Sadd BM, Siva-Jothy MT (2006) Self-harm caused by an insect’s innate immunity. Proceedings of the Royal Society B: Biological Sciences 273, 2571–2574.

Shudo EMI, Iwasa YOH (2001) Inducible Defense against Pathogens and Parasites: Optimal Choice among Multiple Options. Journal of Theoretical Biology 209, 233–247.

Tate AT, Andolfatto P, Demuth JP, Graham AL (2017) The within-host dynamics of infection in trans-generationally primed flour beetles. Molecular Ecology 26, 3794–3807.

Tate AT, Graham AL (2015a) Dynamic Patterns of Parasitism and Immunity across Host Development Influence Optimal Strategies of Resource Allocation. The American Naturalist 186, 495–512.

Tate AT, Graham AL (2015b) Trans-generational priming of resistance in wild flour beetles reflects the primed phenotypes of laboratory populations and is inhibited by co-infection with a common parasite. Functional Ecology 29, 1059–1069.

Tate AT, Graham AL (2017) Dissecting the contributions of time and microbe density to variation in immune gene expression. Proceedings of the Royal Society B: Biological Sciences 284.

Tzou P, Ohresser S, Ferrandon D, et al. (2000) Tissue-Specific Inducible Expression of Antimicrobial Peptide Genes in Drosophila Surface Epithelia. Immunity 13, 737–748.

Ezenwa, V.O Rampal S. Etienne, Gordon Luikart, Albano Beja-Pereira, Anna E. Jolles (2010) Hidden Consequences of Living in a Wormy World: Nematode-Induced Immune Suppression Facilitates Tuberculosis Invasion in African Buffalo. The American Naturalist 176, 613–624.

Westra Edze R, van Houte S, Oyesiku-Blakemore S, et al. (2015) Parasite Exposure Drives Selective Evolution of Constitutive versus Inducible Defense. Current Biology 25, 1043–1049.

Yan G, Stevens L, Goodnight CJ, Schall JJ (1998) Effects of a tapeworm parasite on the competition of Tribolium beetles. Ecology (Washington D C) 79, 1093–1103.

Yokoi K, Koyama H, Ito W, et al. (2012a) Involvement of NF-κB transcription factors in antimicrobial peptide gene induction in the red flour beetle, Tribolium castaneum. Developmental & Comparative Immunology 38, 342–351.

Yokoi K, Koyama H, Minakuchi C, Tanaka T, Miura K (2012b) Antimicrobial peptide gene induction, involvement of Toll and IMD pathways and defense against bacteria in the red flour beetle, Tribolium castaneum. Results in Immunology 2, 72–82.

