## Supplemental Figures and Tables for "Natural variation in the contribution of microbial density to inducible immune dynamics"

**Figure S1.** Correlations in the relative expression levels of immune genes and bacterial density for host species, by site. All values are on a log scale. "Bt" refers to relative bacterial density, and the remainder of the labelled rows are immune genes. The results are shown for single *T. castaneum* and *T. confusum* populations isolated from the same site (column 1: Snavely site; column 2: Green River site) as well as the conglomeration of all populations (right-most column). Colors highlight the strength of the correlation (Pearson correlation; <yellow < 0.5 < blue < 0.65 < light red < 0.85 < dark red < 1). *Cec3* was not assayed for wild-derived beetles due to failure of any primer sets to achieve efficient amplification for all populations.

**Figure S2.** Survival, resistance, and immune gene expression of natural populations during *Photorhabdus luminescens* infection. Survival during acute infection was monitored for 24 hours post infection (**A**; N = 50-60 beetles/population). Relative bacterial density for each individual within each population at 14 hours post infection, as quantified by RT-qPCR, is calculated as the log of the linearized difference between P.lum-specific and host reference gene expression (**B**). The relationship between the log of bacterial density and the log of *attacin*-1 (**C.**) and *defensin-1* (**D.**) gene expression 14 hours after challenge with saline or P.lum infection illustrates variation in the intercept and slope of inducible immune gene expression among populations. Lines represent linear fits for each main variable level as computed by the "lm" function in the geom_smooth algorithm of ggplot2 (R).

**Figure S3.** Correlations between constitutive and inducible immune parameters and phenotypic outcomes of infection with *Bacillus thuringiensis* and *Photorhabdus luminescens*. Quantification of Pearson correlation coefficients (**A;** bottom half = coefficient (significant ones in color), top half = relative magnitude) among phenotypes and immune parameters. Bt density at 8 hours post infection is associated with host mortality (**B**), but P.lum density at 14 hours shows no association with host mortality (**C**). Bt density at 8 hours post infection, given a low initial dose, is highly correlated to bacterial density from higher initial doses (**D**). Bt density at 8 hours post infection is positively correlated to the slope of *defensin-1* expression (**E**). HR = hazard ratio, BLUD = Bt density at death, Bt_dose and Plum_dens = bacterial density at 8 and 14 hours post infection respectively, naïve = expression in uninfected individuals, int = microbe-independent inducible gene expression intercept (saline-injected expression at 8 hours), coef = slope of expression over bacterial density. Att1 = *attacin-1*, Def1 = *defensin-1*, pg = *pgrp-sc2*, ddc = *dopa decarboxylase*. Lines represent linear fits for each main variable level as computed by the "lm" function in the geom_smooth algorithm of ggplot2 (R).

**Figure S4.** Patterns of constitutive and inducible *pgrp-sc2* (**A**), *ddc* (**B**), and *cecropin-3* (**C**) expression by population, and *attacin-1* (**D**) and *cecropin-3* (**E**) expression by RNAi treatment in *T. castaneum*. The relationship between the log of bacterial density and the log of immune gene expression 8 hours after challenge with saline or Bt infection illustrates variation in both the intercept (microbe-independent) and slope (microbe-dependent sensitivity) of inducible immune gene expression among populations. Lines represent linear fits for each main variable level as computed by the "lm" function in the geom_smooth algorithm of ggplot2 (R). Top row: color-coded by population. Bottom row: blue = injected with MalE (non-target) dsRNA, yellow = injected with *imd*-dsRNA prior to infection.

**Figure S5.** Bacterial density and immune gene expression at the time of infection-induced mortality with Bt. Bacterial density as quantified by RT-qPCR in individuals showing moribund behavior (**A**) is not closely associated with the time post infection at which the individual transitions into the moribund group (**B**). Expression of *defensin-1* (**C**) and *ddc* (**D**) as a function of bacterial density shows variation in magnitude among populations. Lines represent linear fits for each main variable level as computed by the "lm" function in the geom_smooth algorithm of ggplot2 (R).

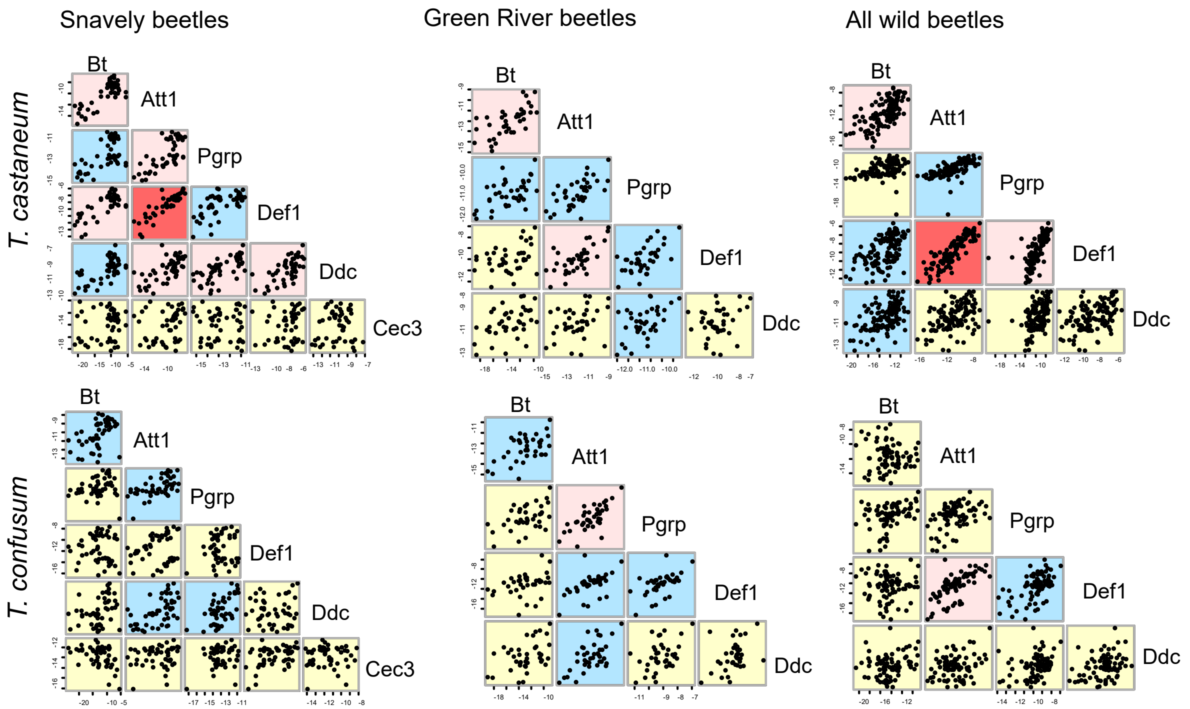

Snavely beetles
Green River beetles
All wild beetles
T. castaneum
T. confusum
Bt
Att1
Pgrp
Def1
Ddc
Cec3

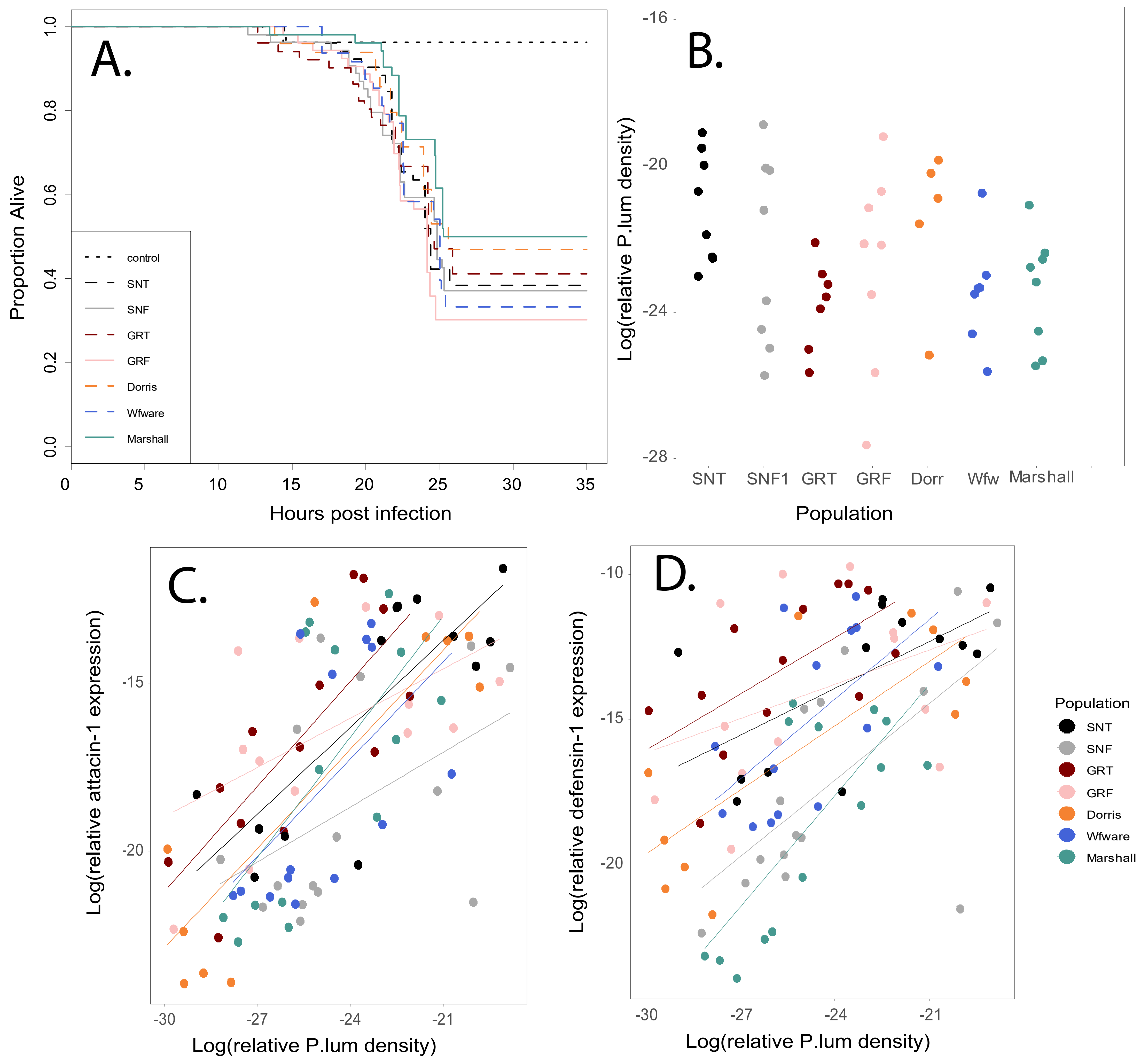

A.
Proportion Alive
Hours post infection
control
SNT
SNF
GRT
GRF
Dorris
Wfware
Marshall
B.
Log(relative P.lum density)
Population
SNT
SNF1
GRT
GRF
Dorr
Wfw
Marshall
C.
Log(relative attacin-1 expression)
Log(relative P.lum density)
D.
Log(relative defensin-1 expression)
Log(relative P.lum density)
Population
SNT
SNF
GRT
GRF
Dorris
Wfware
Marshall

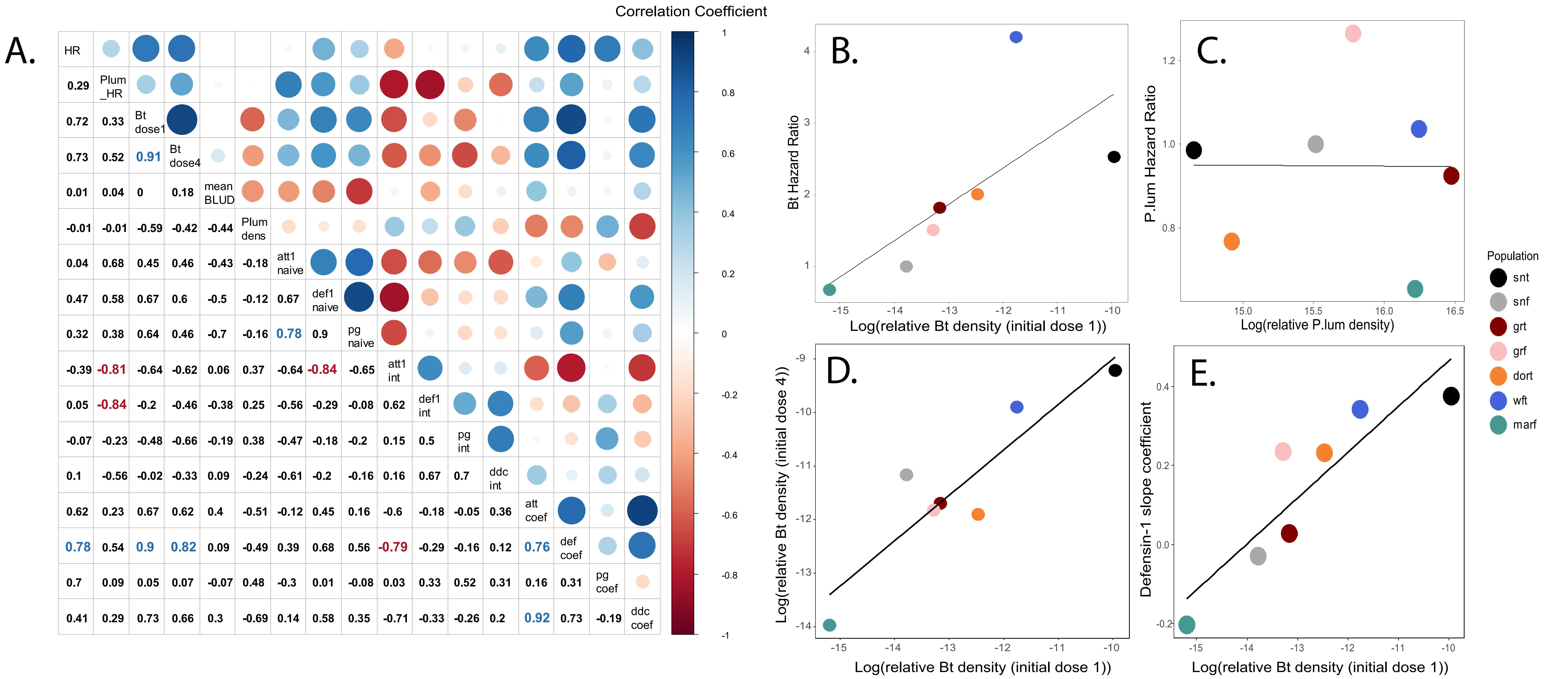
A.
Correlation Coefficient
HR
0.29 Plum_HR
0.72 0.33 Bt dose1
0.73 0.52 0.91 Bt dose4
0.01 0.04 0 0.18 mean BLUD
-0.01 -0.01 -0.59 -0.42 -0.44 Plum dens
0.04 0.68 0.45 0.46 -0.43 -0.18 att1 naive
0.47 0.58 0.67 0.6 -0.5 -0.12 0.67 def1 naive
0.32 0.38 0.64 0.46 -0.7 -0.16 0.78 0.9 pg naive
-0.39 -0.81 -0.64 -0.62 0.06 0.37 -0.64 -0.84 -0.65 att1 int
0.05 -0.84 -0.2 -0.46 -0.38 0.25 -0.56 -0.29 -0.08 0.62 def1 int
-0.07 -0.23 -0.48 -0.66 -0.19 0.38 -0.47 -0.18 -0.2 0.15 0.5 pg int
0.1 -0.56 -0.02 -0.33 0.09 -0.24 -0.61 -0.2 -0.16 0.16 0.67 0.7 ddc int
0.62 0.23 0.67 0.62 0.4 -0.51 -0.12 0.45 0.16 -0.6 -0.18 -0.05 0.36 att coef
0.78 0.54 0.9 0.82 0.09 -0.49 0.39 0.68 0.56 -0.79 -0.29 -0.16 0.12 0.76 def coef
0.7 0.09 0.05 0.07 -0.07 0.48 -0.3 0.01 -0.08 0.03 0.33 0.52 0.31 0.16 0.31 pg coef
0.41 0.29 0.73 0.66 0.3 -0.69 0.14 0.58 0.35 -0.71 -0.33 -0.26 0.2 0.92 0.73 -0.19 ddc coef
1 0.8 0.6 0.4 0.2 0 -0.2 -0.4 -0.6 -0.8 -1
B.
Bt Hazard Ratio
Log(relative Bt density (initial dose 1))
C.
P.lum Hazard Ratio
Log(relative P.lum density)
D.
Log(relative Bt density (initial dose 4))
Log(relative Bt density (initial dose 1))
E.
Defensin-1 slope coefficient
Log(relative Bt density (initial dose 1))
Population
snt
snf
grt
grf
dort
wft
marf

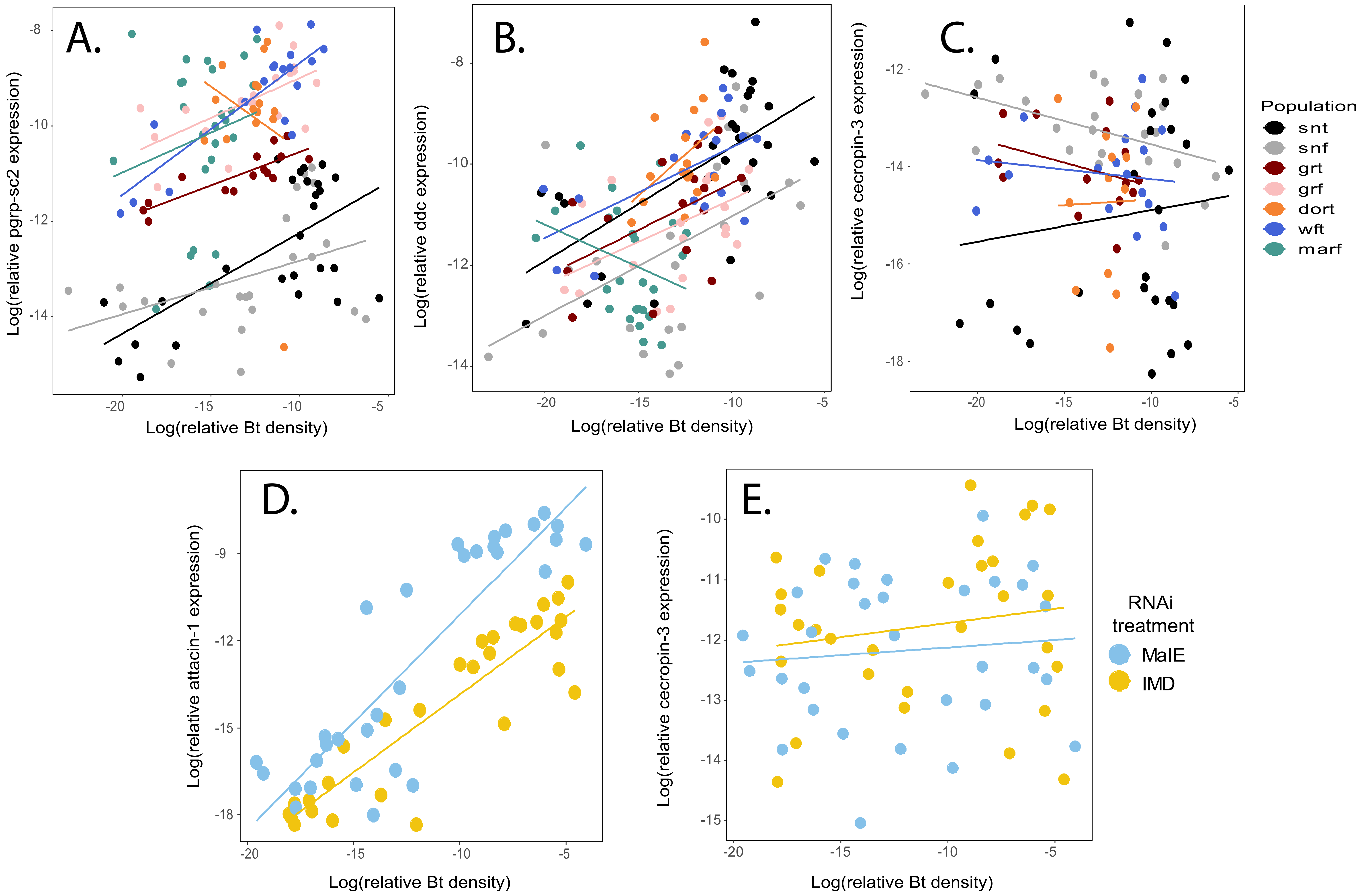

A.
Log(relative pgrp-sc2 expression)
-8
-10
-12
-14
-20
-15
-10
-5
Log(relative Bt density)
B.
Log(relative ddc expression)
-8
-10
-12
-14
-20
-15
-10
-5
Log(relative Bt density)
C.
Log(relative cecropin-3 expression)
-12
-14
-16
-18
-20
-15
-10
-5
Log(relative Bt density)
Population
snt
snf
grt
grf
dort
wft
marf
D.
Log(relative attacin-1 expression)
-9
-12
-15
-18
-20
-15
-10
-5
Log(relative Bt density)
E.
Log(relative cecropin-3 expression)
-10
-11
-12
-13
-14
-15
-20
-15
-10
-5
Log(relative Bt density)
RNAi treatment
MalE
IMD

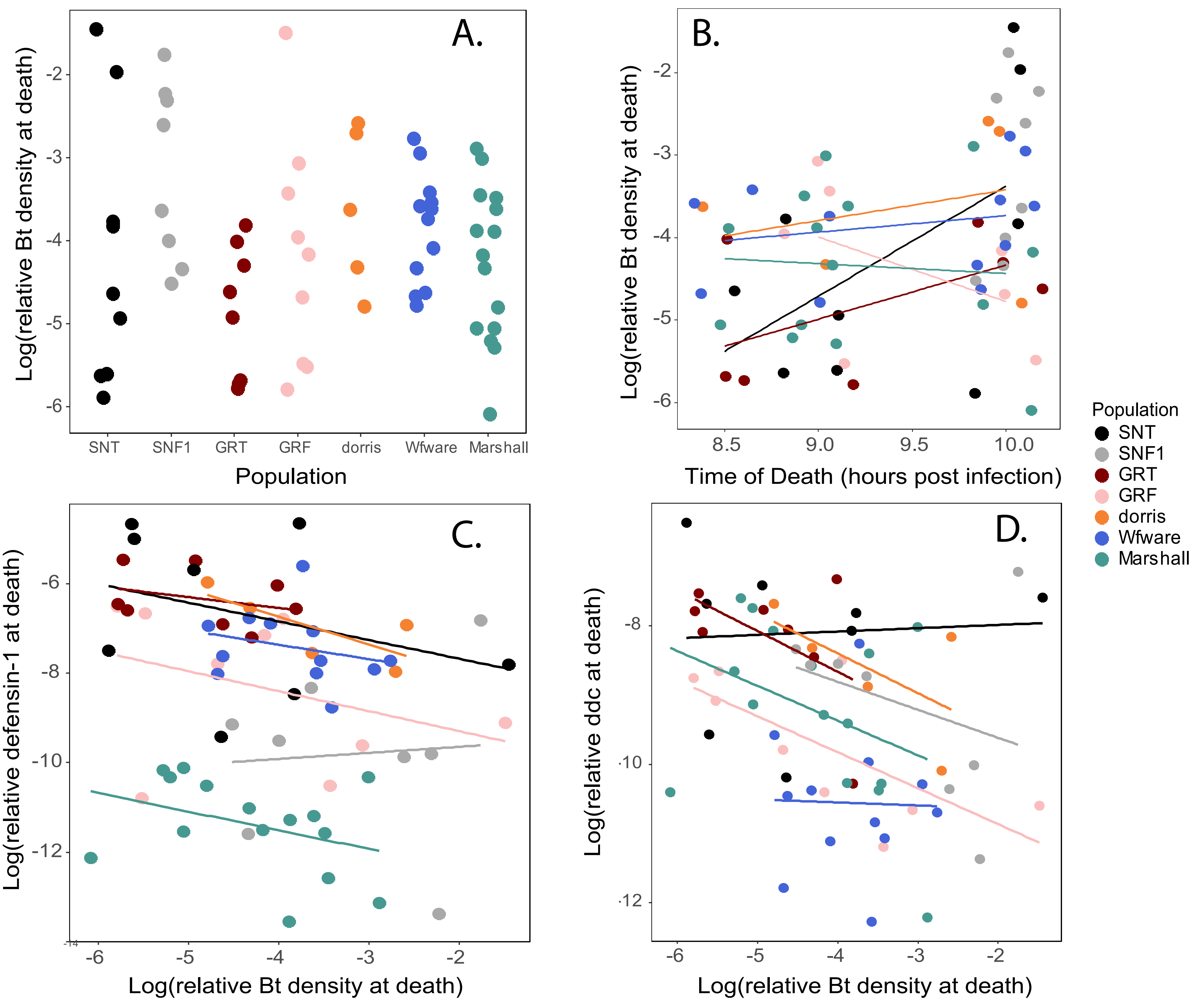
A.
B.
C.
D.
Log(relative Bt density at death)
Population
SNT
SNF1
GRT
GRF
dorris
Wfware
Marshall
Time of Death (hours post infection)
Log(relative defensin-1 at death)
Log(relative ddc at death)
Population
SNT
SNF1
GRT
GRF
dorris
Wfware
Marshall

**Table S1**. Primer sequences used in study, in 5' to 3' order

| Primer Set | Full Name/Function | Forward Oligo Sequence | Reverse Oligo Sequence | AT. (˚C) |
|---|---|---|---|---|
| Att1 | Attacin-1 (IMD AMP) | AAACARTTYCAYCCAAATGG | AGATCKGTWCCRWRTTTCTKGGT | 55 |
| Def1 | Defensin-1 (Toll/IMD AMP) | TTTRYCGTTGCARTAKCCTCC | TCAARSTGAATCATGCCGCWTG | 60 |
| Cec3 | Cecropin-3 (Toll AMP) | AACATGARYACCAAACTTTT | CCAAYTTATMGGCTKTGGWG | 55 |
| PGRP | PGRP-SC2 (IMD recognition) | ACAGTTGGATGCKTTGAAACAGT | AACTSGTYCTGCTCCCTTG | 52 |
| Ddc | Ddc (melanization cascade) | AGAAGTCGTGATGCTKGACT | CTTGRATCACGCCGCC | 55 |
| 18s | RPS18 (ribosomal protein) | CGAAGAGGTCGAGAAAATCG | CGTGGTCTTGGTGTGTTGAC | 58 |
| Bt | Bt 16s rRNA | GACTTTCTGGTCTGTAACTGACA | ACTTCAGCACTAAAGGGCGGA | 55 |
| MalE - dsRNA | Maltose binding (*E. coli* ) | ATTGCTGCTGACGGGGGTTAT | ATGTTCGGCATGATTTCACCTTT | 55 |
| IMD - dsRNA | Imd | CCTCCAAGGGATGAAGTCAA | TTTCCAACAGTGGCACAATC | 55 |
| IMD - qPCR | Imd | AACTGATGCCATACCCAGCC | CAAAAGCAGATGGTCCGCTC | 55 |

**Notes**: AT refers to the primer annealing temp used in a 3 step qPCR sequence. The dsRNA sequences include a 5' end T7 binding sequence, taatacgactcactataggg

**Table S2. Gene expression in naïve and saline-stabbed beetle populations, relative to SNT**

| Gene | Factor | Estimate | Std. Error | t value | Pval |
|---|---|---|---|---|---|
| **Naïve** | | | | | |
| *attacin-1* | Intercept | -19.69 | 0.50 | -39.52 | **<2e-16** |
| | SNF | -1.61 | 0.72 | -2.24 | **0.028** |
| | GRT | -1.02 | 0.70 | -1.45 | 0.15 |
| | GRF | -0.23 | 0.72 | -0.32 | 0.75 |
| | Dorris | -2.48 | 0.74 | -3.37 | **0.0012** |
| | Wfware | -1.78 | 0.70 | -2.52 | **0.014** |
| | Marshall | -1.99 | 0.70 | -2.83 | **0.0059** |
| *defensin-1* | Intercept | -16.60 | 0.50 | -33.50 | **<2e-16** |
| | SNF | -4.19 | 0.72 | -5.86 | **9.90E-08** |
| | GRT | 0.51 | 0.70 | 0.72 | 0.47 |
| | GRF | -0.50 | 0.72 | -0.70 | 0.49 |
| | Dorris | -2.40 | 0.73 | -3.28 | **0.0015** |
| | Wfware | -1.74 | 0.70 | -2.49 | **0.015** |
| | Marshall | -5.75 | 0.70 | -8.20 | **3.31E-12** |
| *pgrp-sc2* | Intercept | -17.05 | 0.46 | -36.99 | **<2e-16** |
| | SNF | -3.69 | 0.67 | -5.54 | **3.75E-07** |
| | GRT | -0.58 | 0.65 | -0.88 | 0.38 |
| | GRF | -1.12 | 0.67 | -1.68 | 0.097 |
| | Dorris | -2.43 | 0.68 | -3.57 | **0.00062** |
| | Wfware | -2.18 | 0.65 | -3.34 | **0.0013** |
| | Marshall | -3.01 | 0.65 | -4.62 | **1.43E-05** |
| **Saline-stabbed, 8 hours post injection** | | | | | |
| *attacin-1* | Intercept | -13.46 | 0.62 | -21.55 | **<2e-16** |
| | SNF | 1.34 | 0.93 | 1.45 | 0.16 |
| | GRT | 0.71 | 0.99 | 0.72 | 0.48 |
| | GRF | -0.75 | 0.93 | -0.81 | 0.43 |
| | Dorris | 0.66 | 0.99 | 0.67 | 0.51 |
| | Wfware | 0.81 | 0.99 | 0.82 | 0.42 |
| | Marshall | 3.21 | 0.93 | 3.47 | **0.0019** |
| *defensin-1* | Intercept | -11.09 | 0.62 | -17.95 | **3.59E-16** |
| | SNF | -1.34 | 0.92 | -1.47 | 0.15 |
| | GRT | 0.91 | 0.98 | 0.93 | 0.36 |
| | GRF | -1.18 | 0.92 | -1.29 | 0.21 |
| | Dorris | 1.38 | 0.98 | 1.41 | 0.17 |
| | Wfware | 0.69 | 0.98 | 0.70 | 0.49 |
| | Marshall | 2.22 | 0.92 | 2.42 | **0.023** |

Notes: linear models (expression ~ population) conducted with "lm" function in R. Expression values are on a log scale.

**Table S3. Population hazard ratios and gene expression means for naïve, saline-injected, and bacterial inf**

| Population | Bt_HR | Plum_HR | Bt_dose1 | Bt_dose4 | meanBLUD | Plum_dens | att1_naive | def1_naive |
|---|---|---|---|---|---|---|---|---|
| snt | 2.53 | 0.98598 | -9.964 | -9.210 | -4.192 | 14.651 | -19.694 | -16.603 |
| snf | 1 | 1 | -13.781 | -11.161 | -3.175 | 15.514 | -21.303 | -20.795 |
| grt | 1.82 | 0.925 | -13.168 | -11.697 | -4.859 | 16.473 | -20.716 | -16.096 |
| grf | 1.51 | 1.266 | -13.287 | -11.821 | -4.177 | 15.777 | -19.922 | -17.104 |
| dort | 2.01 | 0.768 | -12.474 | -11.900 | -3.608 | 14.921 | -22.175 | -19.007 |
| wft | 4.21 | 1.038 | -11.760 | -9.901 | -3.842 | 16.244 | -21.469 | -18.347 |
| marf | 0.67 | 0.654 | -15.192 | -13.965 | -4.283 | 16.218 | -21.688 | -22.349 |

Notes: Hazard ratios (HR) calculated from Cox Proportional Hazards models relative to SNF. All other values
gene expression in saline-stabbed individuals after 8 hours (or 14 hours in Plum experiment (X_14int). X_co
log scale. Only columns through ddc_coef are represented on corr plot

**fected individuals; used in correlation plots**

| pg_naive | att1_int | def1_int | pg_int | ddc_int | att_coef | def_coef | pg_coef | ddc_coef |
|---|---|---|---|---|---|---|---|---|
| -17.053 | -13.459 | -11.085 | -14.461 | -11.687 | 0.377 | 0.376 | 0.111 | 0.330 |
| -20.739 | -12.117 | -12.429 | -13.861 | -12.154 | 0.287 | -0.029 | 0.095 | 0.199 |
| -17.629 | -12.747 | -10.175 | -11.615 | -11.747 | 0.254 | 0.028 | 0.140 | 0.185 |
| -18.171 | -14.209 | -12.268 | -10.182 | -11.624 | 0.250 | 0.235 | 0.167 | 0.169 |
| -19.481 | -12.796 | -9.710 | -9.706 | -10.162 | 0.652 | 0.232 | 0.157 | 0.418 |
| -19.233 | -12.653 | -10.400 | -11.197 | -11.371 | 0.476 | 0.342 | 0.277 | 0.179 |
| -20.067 | -10.249 | -8.865 | -10.165 | -11.105 | -0.127 | -0.202 | 0.168 | -0.167 |

s are population means for 8 hour bacterial density given an initial dose (Bt_doseX) or at 14 hours (Plum

oef represents the slope of expression by Bt density for each gene. Xatdeath is the gene expression in mo

| att1atdeatl | def1atdeat | pgatdeath | ddcatdeatł | att1_14int | def1_14int | pg_14int |
|---|---|---|---|---|---|---|
| -9.566 | -6.696 | -5.131 | -7.981 | -20.128 | -17.846 | -19.422 |
| -11.575 | -9.804 | -6.548 | -9.144 | -22.460 | -18.872 | -21.674 |
| -10.462 | -6.331 | -5.874 | -8.161 | -20.599 | -16.576 | -18.701 |
| -11.198 | -8.321 | -7.413 | -9.736 | -20.310 | -17.304 | -18.106 |
| -11.334 | -6.982 | -7.139 | -8.626 | -19.980 | -17.184 | -18.474 |
| -12.063 | -7.412 | -7.802 | -10.558 | -20.243 | -16.765 | -18.575 |
| -11.959 | -11.395 | -6.756 | -9.229 | -21.366 | -21.165 | -19.708 |

_dens). Gene_naive is average expression in unstabbed individuals. Gene_int is
ɔribund individuals. Note that bacterial densities and gene expressions are on a

**Table S4. Bacterial load at time of death, correlations with immune gene expression, and population**

| Analysis | Factor | Df | Sum Sq | Mean Sq | Fval | P val |
|---|---|---|---|---|---|---|
| **Bacterial Load at time of death (BLUD)** | Population | 6 | 15.13 | 2.521 | 2.33 | **0.0442** |
| | Time of death | 1 | 2.26 | 2.262 | 2.09 | 0.1537 |
| | Residuals | 57 | 61.69 | 1.082 | | |
| **Gene Expression** | | | | | | |
| *attacin-1* | Bt density | 1 | 31.87 | 31.87 | 13.233 | **0.000641** |
| | population | 6 | 33.27 | 5.55 | 2.303 | **0.048221** |
| | Bt*population | 6 | 11.18 | 1.86 | 0.774 | 0.594316 |
| | Residuals | 51 | 122.82 | 2.41 | | |
| *defensin-1* | Bt density | 1 | 16.02 | 16.02 | 8.675 | **0.00485** |
| | population | 6 | 222.23 | 37.04 | 20.06 | **7.03E-12** |
| | Bt*population | 6 | 2.49 | 0.42 | 0.225 | 0.96688 |
| | Residuals | 51 | 94.17 | 1.85 | | |
| *pgrp-sc2* | Bt density | 1 | 32.42 | 32.42 | 17.86 | **9.84E-05** |
| | population | 6 | 20.11 | 3.35 | 1.846 | 0.109 |
| | Bt*population | 6 | 5.19 | 0.87 | 0.477 | 0.823 |
| | Residuals | 51 | 92.59 | 1.82 | | |
| *ddc* | Bt density | 1 | 13.96 | 13.958 | 11.238 | **0.001516** |
| | population | 6 | 38.01 | 6.335 | 5.1 | **3.62E-04** |
| | Bt*population | 6 | 4.01 | 0.669 | 0.538 | 0.776533 |
| | Residuals | 51 | 63.34 | 1.242 | | |

Notes: ANOVAs (expression ~ population*BLUD) conducted with "aov" function in R.

**variation**
